# Direct capsid labeling of infectious HIV-1 by genetic code expansion allows detection of largely complete nuclear capsids and suggests nuclear entry of HIV-1 complexes via common routes

**DOI:** 10.1101/2021.09.14.460218

**Authors:** Sandra Schifferdecker, Vojtech Zila, Thorsten G. Müller, Volkan Sakin, Maria Anders-Össwein, Vibor Laketa, Hans-Georg Kräusslich, Barbara Müller

## Abstract

The cone-shaped mature HIV-1 capsid is the main orchestrator of early viral replication. After cytosolic entry, it transports the viral replication complex along microtubules towards the nucleus. While it was initially believed that the reverse transcribed genome is released from the capsid in the cytosol, recent observations indicate that a high amount of capsid protein (CA) remains associated with subviral complexes during import through the nuclear pore complex (NPC). Observation of post-entry events *via* microscopic detection of HIV-1 CA is challenging, since epitope shielding limits immunodetection and the genetic fragility of CA hampers direct labeling approaches. Here, we present a minimally invasive strategy based on genetic code expansion and click chemistry that allows for site-directed fluorescent labeling of HIV-1 CA, while retaining virus morphology and infectivity. Thereby, we could directly visualize virions and subviral complexes using advanced microscopy, including nanoscopy and correlative imaging. Quantification of signal intensities of subviral complexes revealed an amount of CA associated with nuclear complexes in HeLa-derived cells and primary T cells consistent with a complete capsid and showed that treatment with the small molecule inhibitor PF74 did not result in capsid dissociation from nuclear complexes. Cone-shaped objects detected in the nucleus by electron tomography were clearly identified as capsid-derived structures by correlative microscopy. High-resolution imaging revealed dose-dependent clustering of nuclear capsids, suggesting that incoming particles may follow common entry routes.

## Introduction

The cone-shaped capsid that encases the viral RNA genome and replication proteins is a characteristic feature of infectious human immunodeficiency virus type 1 (HIV-1) particles. Data obtained by many research groups over the past decade have revised our understanding of the role of the mature capsid in HIV-1 replication, placing this structure at the center stage of post-entry replication steps (reviewed in (Aiken and Rousso, 2021; Engelman, 2021; James, 2019; Zila et al., 2021b). Upon fusion of the virion envelope with the cell membrane, the capsid, which consists of ∼1,200-1,500 monomers of the capsid protein CA (Briggs et al., 2003), is released into the cytosol. It then usurps host cell factors to traffic towards the nucleus. Reverse transcription of the viral RNA into dsDNA is initiated during passage of the subviral structure through the cytosol. Following import into the nucleus, the viral dsDNA is covalently integrated into the host cell genome by the viral integrase (IN). Prior to integration, the surrounding capsid shell needs to release the dsDNA in a process termed uncoating. The precise mechanisms, location, and timing of HIV-1 capsid uncoating are still under investigation.

Initially, the HIV-1 capsid was presumed to rapidly dissociate upon cell entry, based on little or no CA detected associated with isolated post-entry complexes (reviewed in (Campbell and Hope, 2015)). Rapid or gradual disassembly in the cytosol was also supported by several studies that applied fluorescence imaging to analyze subviral complexes in infected cells (e.g., (Hulme et al., 2011; Mamede et al., 2017; Xu et al., 2013). However, HIV-1 CA or the capsid lattice was found to directly engage in interactions with various host factors involved in post-entry replication steps. Among these capsid-interacting host factors are not only cytosolic proteins, including proteins involved in microtubular transport, but also nucleoporins and even the nuclear protein cleavage and polyadenylation specific factor 6 (CPSF6) (reviewed in (Engelman, 2021; Naghavi, 2021; Saito and Yamashita, 2021)). These findings implied involvement of at least a partial lattice structure in later stages of post-entry replication. Furthermore, increasing evidence from imaging-based analyses argued for capsid uncoating at or close to the nuclear pore (Burdick et al., 2017; Francis et al., 2020a; Francis et al., 2016; Francis and Melikyan, 2018), or even indicated passage of (nearly) intact capsids through nuclear pores (Burdick et al., 2020; Li et al., 2021; Muller et al., 2021; Zila et al., 2021a). The recent detection of cone-shaped objects in the nuclear pore channel and inside the nucleus by correlative light and electron microscopy (CLEM) (Zila et al., 2021a), and intranuclear separation of CA or IN from reverse transcribed dsDNA (Muller et al., 2021) also supported the model that the nucleus is the site of HIV-1 uncoating (Muller et al., 2022).

One explanation for apparent discrepancies between different studies are the methods that have been used for CA detection in fluorescence microscopy. Since the modification of CA by genetic labeling strategies proved to be challenging, most studies applied immunofluorescence (IF) staining or indirect labeling through a capsid binding protein (e.g., (Burdick et al., 2017; Francis et al., 2016; Hulme et al., 2015; Mamede et al., 2017; Peng et al., 2014; Zila et al., 2019)). A limitation of IF is that staining efficiency may vary substantially depending on the antibody and detection conditions used, as well as on differential exposure or shielding of epitopes due to conformational changes or different intracellular environments. We could indeed show previously that immunostaining efficiency of CA in the nucleus of host cells strongly depends on cell type and experimental conditions (Muller et al., 2021). Furthermore, IF is incompatible with live cell analyses. Infectious HIV-1 derivatives carrying fluorescent CA would resolve these limitations and allow the direct observation of entering capsids with quantitative analyses.

Direct genetic labeling of viral capsid proteins is challenging, however. Capsid proteins are generally small proteins that need to assemble into ordered multimeric lattices. The resulting assemblies must be stable during virus formation and transmission to a new target cell, but also ready to disassemble in the newly infected cell, requiring structural flexibility of the protomers. Beyond protein-protein interactions involved in capsid assembly itself, capsid proteins generally undergo crucial interactions with other components of the virion, e.g., the viral genome. Finally, the capsid surface represents an essential contact interface between virus and host cell in the early phase of infection, mediating cell entry in the case of non-enveloped viruses or interacting with critical host cell dependency or restriction factors in the case of enveloped viruses. Consequently, a large proportion of the surface exposed amino acids of a viral capsid protein is involved in intermolecular contacts that are crucial for virus replication, which renders these proteins highly susceptible to genetic modification. Fusion of a capsid protein to a relatively large genetic label, e.g., green fluorescent protein (GFP) or other fluorescent proteins, is thus generally prone to severely affect virus infectivity.

These considerations also apply to HIV-1 CA. The protein is encoded as a subdomain of the structural polyprotein Gag, from which it is released by the viral protease (PR) concomitant with virus budding to allow for formation of the mature capsid. With a molecular mass of ∼24 kDa, mature CA is of a similar size as GFP. CA hexamers are the core structural elements of the immature Gag shell forming the nascent virus bud in HIV-1 producing cells. Hexamers with a different conformation, together with 12 CA pentamers that allow for bending and curvature, are the building blocks of the mature capsid. CA pentamers, immature and mature hexamers employ different protein-protein interfaces; together, these interfaces involve most of the exposed surface of the CA monomer (reviewed in (Mattei et al., 2016)). Accordingly, scanning mutagenesis analyses found HIV-1 CA to be highly genetically fragile (Rihn et al., 2013; von Schwedler et al., 2003), with up to 89% of single amino acid exchanges tested abolishing or severely affecting virus replication (Rihn et al., 2013). It is thus not surprising that the introduction of genetically encoded labels - GFP or even a small peptide tag - at various positions within HIV-1 CA have resulted in loss or severe reduction of infectivity. Complementation with wild-type (wt) virus, from at least equimolar amounts of wt CA to a substantial molar excess, was essential to restore virus infectivity (Burdick et al., 2020; Campbell et al., 2008; Pereira et al., 2011; Zurnic Bonisch et al., 2020). While the use of wt complemented particles can be sufficient for fluorescent labeling, it is unclear whether the modified CA molecules are an integral part of the mature CA lattice. Only approximately half of the CA molecules present inside the virion are eventually used to form the mature capsid (Briggs et al., 2004; Lanman et al., 2004), and CA fusion proteins may be preferentially excluded or less stably integrated into the mature lattice.

We therefore established and applied a minimal invasive labeling strategy for HIV-1 CA based on genetic code expansion (GCE) and click labeling. This method involves the exchange of a selected amino acid residue in the protein of interest with a non-canonical amino acid (ncAA) carrying a highly reactive bio-orthogonal functional group by a process termed amber suppression (Figure 1a). This residue is subsequently covalently coupled to a fluorophore functionalized with a cognate reaction partner (Figure 1a, b; reviewed in e.g. (Lang and Chin, 2014; Muller et al., 2019; Nikic and Lemke, 2015)). Using this approach, we generated a CA-labeled HIV-1 derivative that largely retained infectivity. In contrast to previous approaches for direct CA labeling, our minimally modified derivative did not require complementation with wt virus. Direct labeling with a bright and photostable chemical dye allows the application of various imaging methods, i.e., live-cell imaging, super-resolution nanoscopy, or CLEM. The click labeled virus variant thus enabled us to directly assess the amount of CA associated with entering subviral complexes outside and within the nucleus of infected HeLa-derived cells and primary CD4^+^ T-cells, to visualize CA containing structures in the nucleus by nanoscopy and correlative microscopy and to study the effect of the CA-binding drug PF74 on the nuclear complexes.

**Figure 1.**
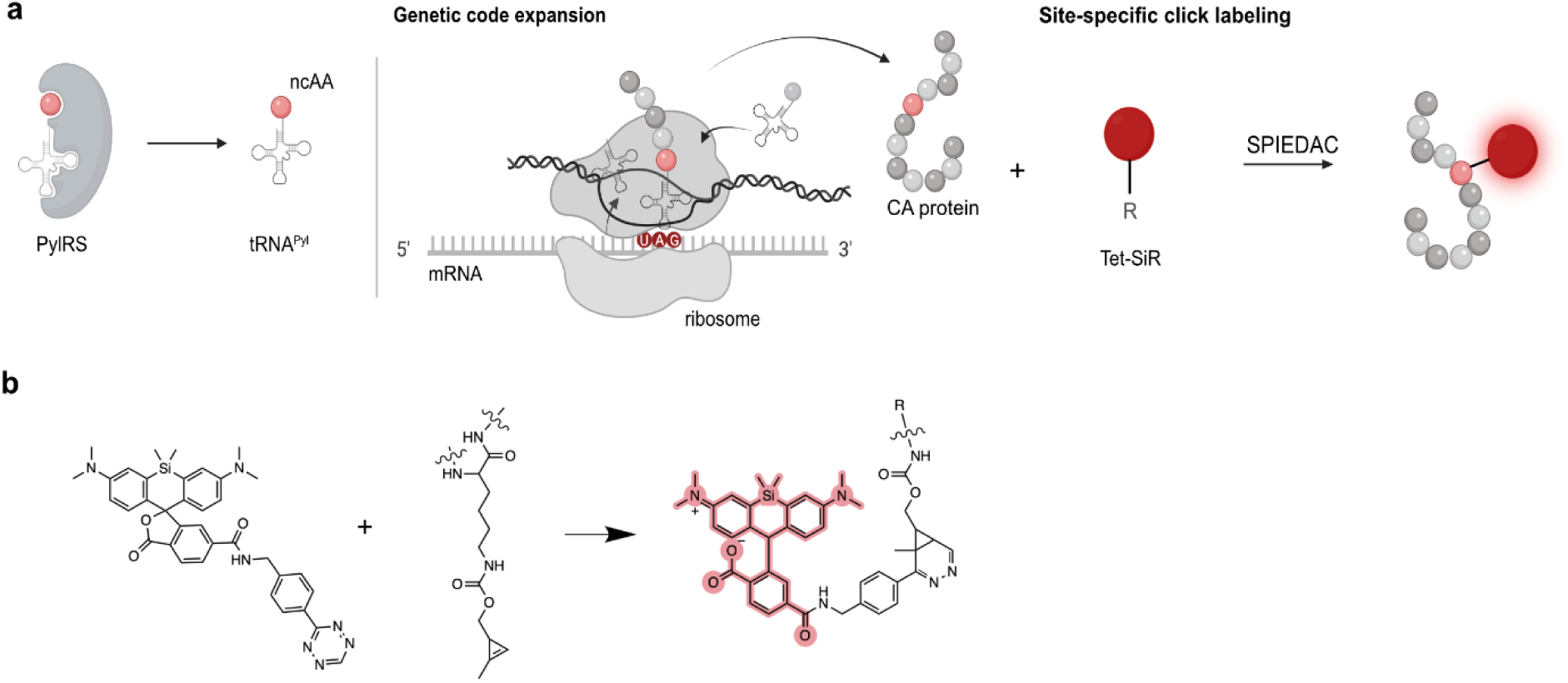
GCE and click labeling of HIV-1 CA. **(a)** Experimental scheme for GCE and click-labeling. The system used here requires the introduction of an amber stop codon (UAG) at a specific site into the CA coding sequence. A genetically engineered bio-orthogonal tRNA / aminoacyl-tRNA synthetase pair mediates incorporation of a non-canonical amino acid (ncAA) at the chosen position. In a second step, a highly reactive group of the ncAA is covalently linked to a fluorophore carrying a cognate reactive group (e.g., a tetrazine group reacting with a cyclopropene group at the ncAA via strain-promoted inverse electron-demand Diels-Alder cycloaddition (SPIEDAC)). Image created with BioRender.com. **(b)** The tetrazine-derivative of silicon rhodamine (Tet-SiR) reacts with the strained alkene of the ncAA cyclopropene-L-lysine (CpK) via SPIEDAC. The open SiR conformation results in fluorescence (highlighted in red).

## Results

### Generation of an HIV-1 variant carrying a bio-orthogonal amino acid within CA

To allow for minimally invasive labeling of HIV-1 CA by GCE (Figure 1), we introduced an amber stop codon at a position of interest into the CA coding sequence within the *gag* open reading frame of the proviral plasmid pNLC4-3 (Bohne and Krausslich, 2004). In order to avoid GCE modification of the accessory viral protein R (Vpr), which is incorporated into the virion in high amounts (Muller et al., 2000), we first exchanged the amber stop codon of *vpr* to an opal codon (TGA), resulting in plasmid pNLC4-3*. Although this mutation did not alter the coding sequence of viral proteins or virion infectivity (Figure S1), the corresponding virus was termed HIV-1* to indicate the modification. Since neither the efficiency of amber suppression in a given sequence context in eukaryotic cells, nor the effect of ncAA incorporation on viral functionality can be predicted with certainty, we tested a panel of 18 amber mutations at sites located towards the outer surface of the capsid lattice for suppression efficiency and virus infectivity (Schifferdecker, Sakin et al., in preparation). Based on a comparison of Gag expression levels and viral infectivity upon ncAA incorporation, we selected a virus variant in which residue alanine 14 in CA was replaced by a non-canonical amino acid (HIV-1*CA14^ncAA^) for further analyses.

For virus preparation, HEK293T cells were co-transfected with the respective mutant proviral plasmid and pNESPylRS-eRF1dn-tRNA. The latter plasmid encodes for a complete amber suppression system, consisting of modified tRNA, a cognate genetically engineered pyrrolysine aminoacyl-tRNA synthetase (Nikic et al., 2016), and a dominant-negative version of the eukaryotic release factor eRF1 that improves amber suppression efficiency in eukaryotic cells (Schmied et al., 2014). To produce functionalized virus particles, cells were grown in the presence of the small ncAA cyclopropene lysine (CpK). While truncation of Gag at position 14 of CA would prevent virus formation, incorporation of CpK by amber suppression should result in the expression of full-length Gag and thereby promote HIV-1 particle assembly. Immunoblot analysis of cell lysates indeed demonstrated the presence of full-length Gag polyprotein precursor when HIV-1*CA14^TAG^ expressing cells were grown in the presence of CpK, whereas full-length Gag was not detected when CpK was omitted from the growth medium (Figure S2a). Thin-section electron microscopy (EM) revealed late budding sites and immature-as well as mature-like virions at the plasma membrane and in the vicinity of HIV-1*CA14^TAG^ expressing cells, that were morphologically indistinguishable from typical HIV-1 wild-type (wt) budding sites and virions (Figure 2a). Virus release efficiency from transfected cells, as estimated by immunoblot analysis of cell and particle lysates, was comparable for wt and HIV-1*CA14^CpK^ (Figure S2b). We concluded that Gag expression of HIV-1*CA14^TAG^ is ncAA dependent and the modified CA domain is competent for immature and mature lattice assembly.

**Figure 2.**
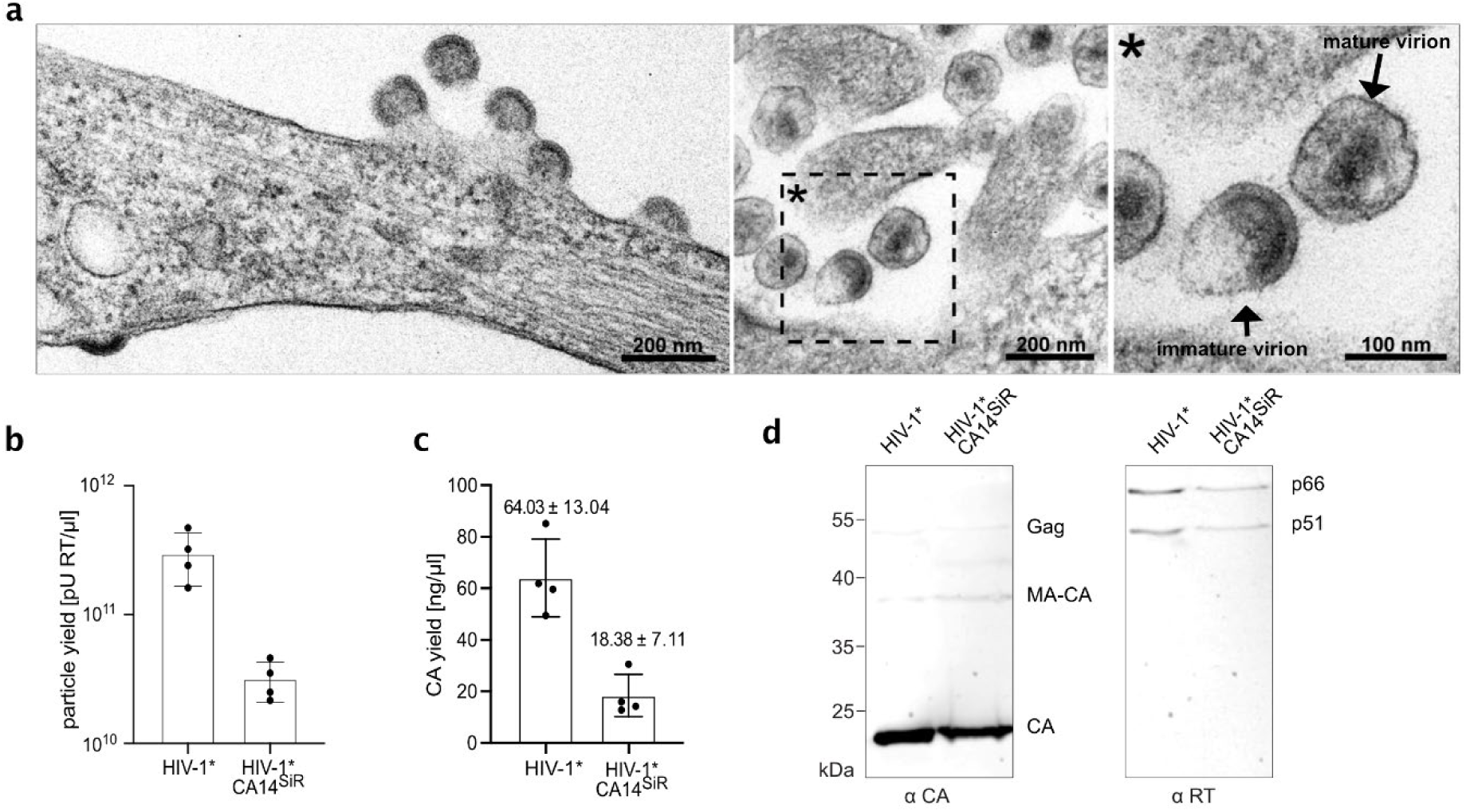
Production and characterization of click labeled HIV-1 (HIV-1*CA14^SiR^). **(a)** Morphology of HIV-1*CA14^ncAA^ assembly sites and particles. HEK293T cells were co-transfected with pNLC4-3*CA14^TAG^ and pNESPylRS-eRF1dn-tRNA and grown in the presence of 500 µM CpK. At 44 h p.t., cells were fixed, embedded, and analyzed by thin-section EM as described in materials and methods. **(b,c)** Virus production. Click labeled particles were prepared from the supernatant of HEK293T cells co-transfected with either pNLC4-3* or pNLC4-3*CA14^TAG^ and pNESPylRS-eRF1dn-tRNA and grown in the presence of 500 µM CpK as described in materials and methods. Particle yield in the final preparations was determined via quantitation of RT activity (SG-PERT assay; (Pizzato et al., 2009)) (b) and by determination of CA amounts using quantitative immunoblot as described in materials and methods (c). The graphs show mean values and SD from four independent experiments. **(d)** Immunoblot analysis of virus preparations. 5 µl HIV-1* and HIV-1*CA14^SiR^ particle lysates were separated by SDS-PAGE, and proteins were transferred to nitrocellulose membranes by semi-dry blotting. Viral proteins were detected using the indicated polyclonal antisera. Bound antibodies were detected by quantitative immunofluorescence with a Li-COR CLx infrared scanner, using secondary antibodies and protocols according to the manufacturer’s instructions. Positions corresponding to Gag/Gag-Pol and processing products are indicated to the right.

### Characterization of click labeled HIV-1 virions

We next prepared virus particles from the supernatant of HIV-1*CA14^ncAA^ producing cells and subjected them to click labeling using the membrane-permeable dye silicon rhodamine tetrazine (SiR-Tet; (Lukinavicius et al., 2013); Figure 1b), generating HIV-1*CA14^SiR^. As a control, HIV-1* wt particles were prepared under amber suppression conditions and stained in parallel. Consistent with the detection of viral assembly sites and particles in electron micrographs of transfected cells (Figure 2a), virus was recovered from the tissue culture supernatant of HIV-1*CA14^ncAA^ expressing cells. Particle yields were moderately reduced compared to the HIV-1* wt control, in line with the fact that amber suppression is usually incomplete in eukaryotic cells (optimal ncAA incorporation efficiencies in the range of ∼25-50 %; e.g., (Sakin et al., 2017; Schmied et al., 2014). On average, we obtained 5-10-fold lower yields for HIV-1*CA14^SiR^ compared to HIV-1* as determined by RT activity (Figure 2b) and CA content (Figure 2c) of particle preparations. Comparison of stained and unstained preparations demonstrated that particle yield, assessed by RT activity of virus preparations, was not affected by the click-labeling procedure (Figure S3). Consistent with the observation of morphologically mature particles by EM, click labeled particles displayed regular Gag and GagPol processing products (Figure 2d), with clear bands for mature CA (p24) and mature RT heterodimer (p51, p66).

### Fluorescence labeling and infectivity of click labeled virions

In order to determine fluorescence staining specificity, particle lysates of HIV-1* and HIV-1*CA14^CpK^ stained with Tet-SiR were analyzed by SDS-PAGE followed by in-gel fluorescence measurement (Figure 3a). The resulting image revealed a distinct SiR labeled band corresponding to a mass of approximately 24 kDa and weak bands corresponding in size to CA containing precursors in the case of HIV-1*CA14^SiR^, whereas fluorescent protein bands were undetectable for the HIV-1* control (Figure 3a, bottom panel). Subsequent immunoblotting of the scanned gel using antiserum raised against CA ensured that similar amounts of virus particles had been loaded for wt and the CA14 variant (Figure 3a, top panel). These results indicated specific GCE-dependent labeling of CA *via* amber suppression at position 14 of HIV-1 CA.

**Figure 3.**
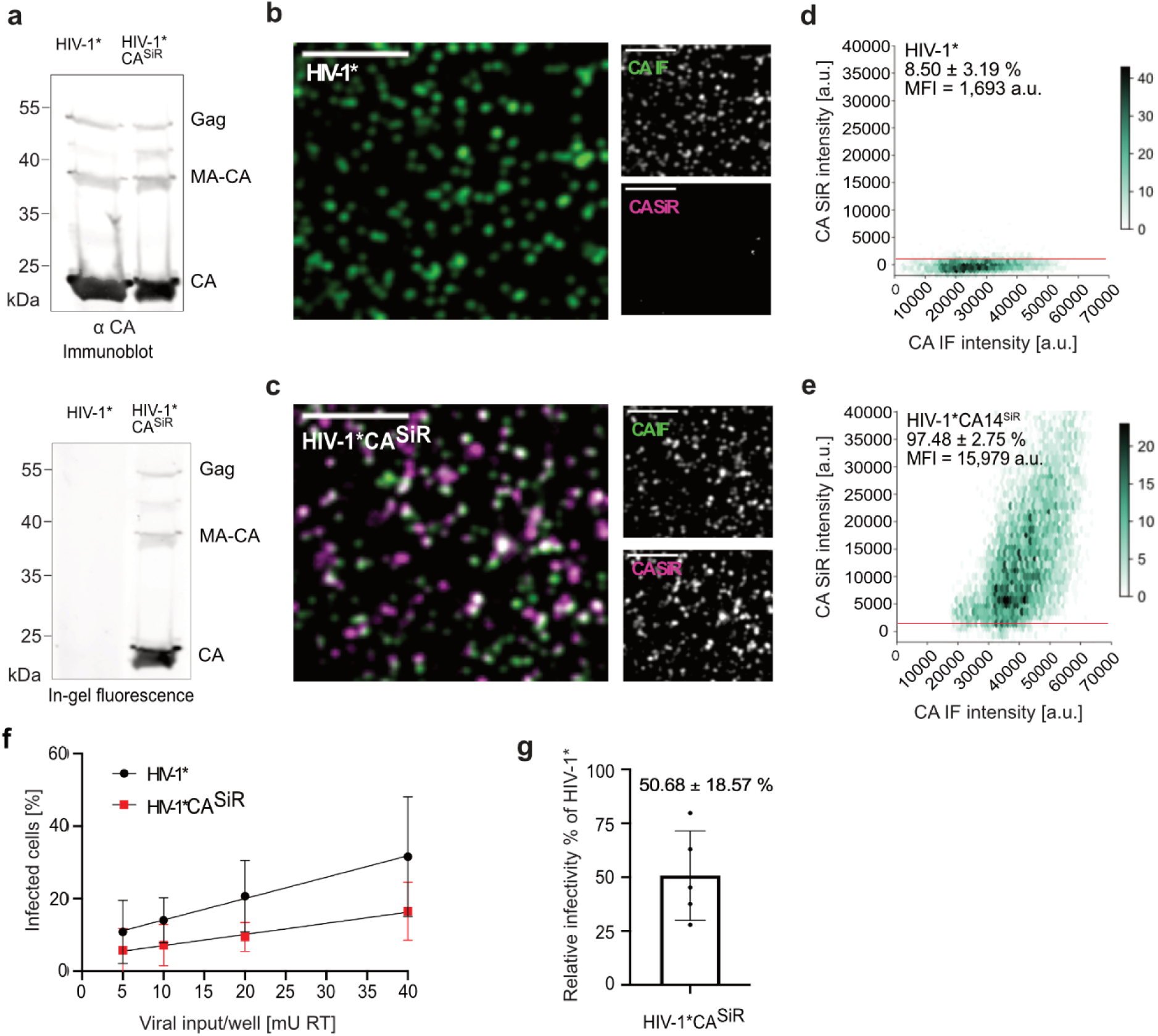
Characterization of CA click labeled particles. **(a)** Specific fluorescent labeling of CA14^CpK^. Immunoblot analysis of virus preparations. 20 µl HIV-1* and HIV-1*CA14^SiR^ particle lysates were separated by SDS-PAGE. In-gel fluorescence (bottom panel) was detected by scanning the gel on a Li-COR Clx infrared scanner set at an emission wavelength of 700 nm. Subsequently immunoblot analysis (top panel) was performed using the previously scanned gel. Proteins were transferred from the gel to a nitrocellulose membrane by semi-dry blotting. Viral proteins were detected using polyclonal antiserum raised against recombinant CA. Bound antibodies were detected by quantitative immunofluorescence with a Li-COR CLx infrared scanner, using secondary antibody and protocol according to the manufacturer’s instructions. Please note that at the high amounts of particles loaded for efficient in-gel fluorescence detection the immunoblot detection is not within the linear range of the method, so that contaminating precursor bands appear overrepresented. **(b-e)** Analysis of labeling efficiency. Particles harvested from the supernatant of virus producing HEK239T cells were subjected to click labeling. Particles were then immobilized on PEI coated chamber slides, fixed, and permeabilized. Particles were immunostained using antiserum raised against HIV-1 CA, and specimens were imaged by SDCM (b,c). Scale bars 5 μm. **(d,e)** Hexabin plots of particles detected in (b,c). Mean intensities of CA(SiR) are plotted against mean intensity CA(IF) for HIV-1* and HIV-1*CA14^SiR^. The color intensity of the hexagons corresponds to the number of particles displaying the indicated intensity values. The graphs represent pooled data from 12 fields of view from three independent virus preparations. The red line indicates the threshold t=1,000. **(f)** Infectivity of click labeled particles. The indicated virus particles were prepared as in (b-e) and subjected to click labeling. Particle yield was assessed by RT activity assay (Pizzato et al., 2009), and samples were titrated on TZM-bl indicator cells seeded in 15-well ibidi µ-slide angiogenesis dishes. 50 µM T-20 was added at 6 h p.i. to prevent second-round infection in the case of HIV-1*. Cells were fixed, permeabilized, and immunostained using a polyclonal rabbit antiserum raised against recombinant HIV-1 MA at 48 h p.i. Samples were imaged by SDCM. The percentage of infected cells was determined using Fiji software. The graphs show mean values and SD from six independent infection experiments using five independent particle preparations (n=5,700-7,700 cells were counted per condition). Lines represent linear regression based on the mean values. **(g)** Relative infectivity of a virus preparation (% infected cells/mU RT) was determined as in (f), and the values obtained for HIV-1*CA14^SiR^ were normalized to the value obtained for HIV-1* virus in the same experiment. All cells counted in (f) were used for quantification. The graph represents the mean value and SD from six independent experiments.

To test efficiency of SiR staining, labeled particles adhered to a glass chamber slide were fixed, permeabilized, immunostained with antiserum raised against HIV-1 CA to validate that detected signals corresponded to virus particles and imaged by confocal microscopy (Figure 3b, c). Confocal micrographs were recorded in the channels corresponding to the CA (IF) stain (green) and direct CA labeling with SiR (magenta). Regions of interest (ROIs) corresponding to the position of virus particles were defined based on CA(IF) signals. Measurement of SiR fluorescence intensities in these ROIs revealed only weak background staining in the case of HIV-1* (Figure 3b). In contrast, distinct SiR signals co-localizing with CA(IF) punctae were detected for HIV-1*CA14^SiR^ (Figure 3c). Quantitative analyses of images from multiple independent experiments confirmed this visual impression (Figure 3d, e). Only ∼8.5% of HIV-1* particles were classified as SiR positive, with fluorescence intensities only slightly above the background level (∼1,000 a.u.; Figure 3d). In contrast, >95% HIV-1*CA14^SiR^ particles displayed clear SiR staining, with a mean fluorescence intensity of ∼15,000 a.u. (Figure 3e). Variation in SiR fluorescence intensities between individual particles is expected, since particle size and CA content of HIV-1 virions varies, with ∼1,700-3,100 CA monomers estimated per particle (Carlson et al., 2008). Beyond that, the range of SiR signal intensities observed also indicated a range of click labeling efficiencies. Despite variable staining intensities within the preparation, the vast majority of HIV-1*CA14^CpK^ particles could be efficiently click labeled with SiR, attaining fluorescence intensities suitable for fluorescence microscopy of infected cells.

To test the effect of introducing a synthetic fluorophore at position 14 on CA functionality, the infectivity of click labeled particles was assessed by titration of labeled particles on TZM-bl cells, followed by immunostaining against the HIV-1 matrix protein (MA) to identify infected cells. As shown in Figures 3f and g, relative infectivity of HIV-1*CA14^SiR^ was only mildly reduced by an average of ∼2-fold compared to HIV-1*. This moderate reduction in infectivity represented a substantial improvement compared to previous genetically labeled derivatives in the absence of complementation (Burdick et al., 2020; Campbell et al., 2008; Pereira et al., 2011; Zurnic Bonisch et al., 2020). Thus, minimal invasive labeling by GCE allows direct labeling of HIV-1 CA without requiring complementation with wt virus.

### Detection of click labeled HIV-1 in infected cells

Having established a suitable labeling strategy, we used labeled particles to infect and image target cells. Initial experiments were performed in the model cell line HeLa TZM-bl. Cells infected with HIV-1*CA14^SiR^ at a multiplicity of infection (MOI) of ∼0.8 were fixed at 18 h post infection (p.i.). Immunostaining with antiserum against CA was performed under conditions that allow for immunodetection of cytosolic and nuclear complexes (Muller et al., 2021) to test whether detected SiR signals corresponded to HIV-1 particles. Labeled particles could be visualized by spinning disc confocal microscopy (SDCM) in the cellular environment (Figure 4a, Figure S4 and S5). Confocal images revealed punctate SiR signals in the cytosol, close to the nuclear envelope and within the nucleus of infected cells. Co-localization with CA(IF) staining confirmed that these signals represented entering viral structures (Figure 4a, Figure S4 and S5).

**Figure 4.**
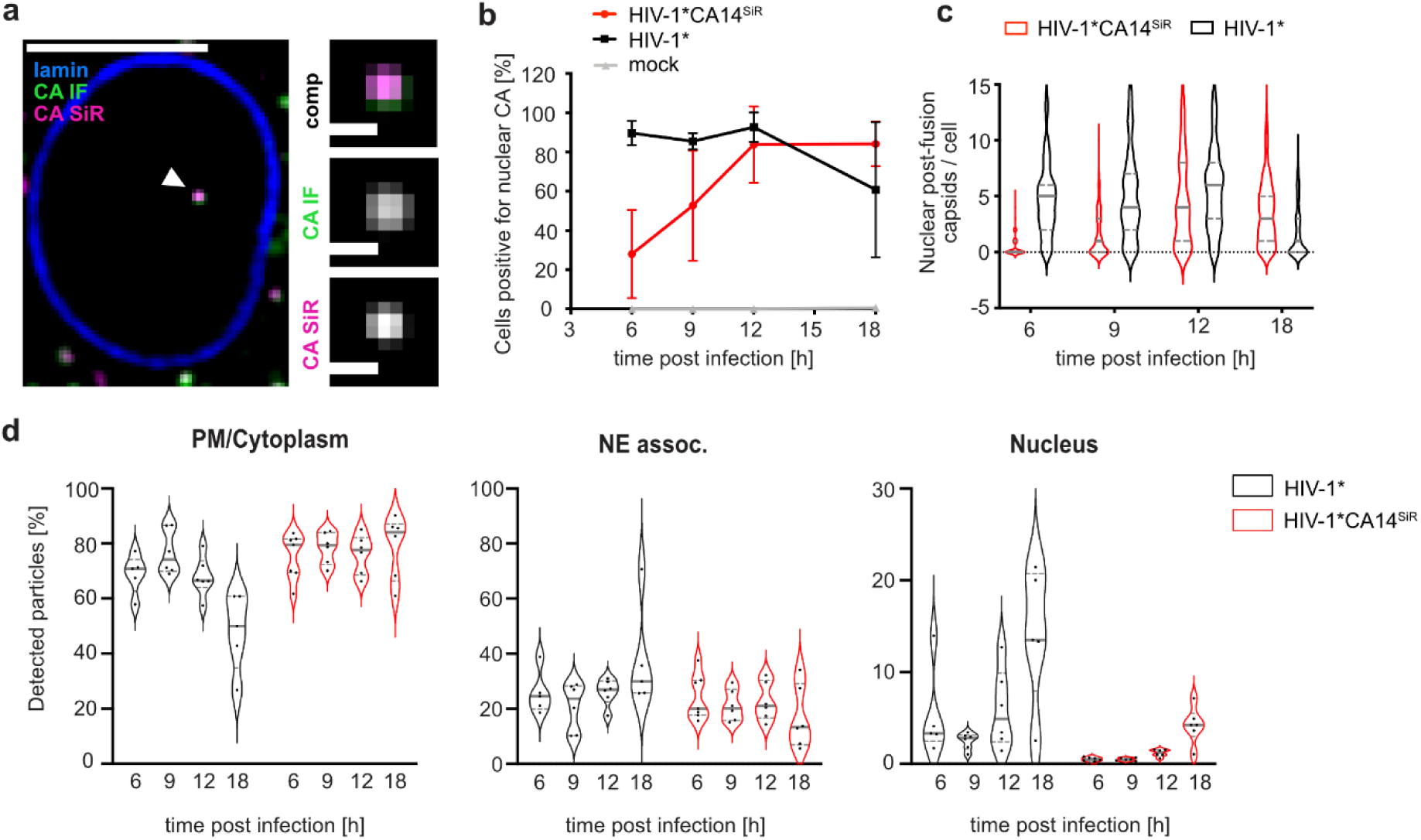
Detection of CA in the nucleus of infected HeLa-derived cells. TZM-bl cells were infected with HIV-1* or HIV-1* CA14^SiR^ particles (∼MOI 0.8), treated with 15 µM PF74 for 1 h before fixation, fixed at 6, 9, 12 and 18 h p.i. and imaged by SDCM. **(a)** Single z slice of a representative cell infected with HIV-1* CA14^SiR^ at 18 h p.i. and one enlarged z slice are shown. Scale bars: 10 µm (cell) and 1 µm (enlargement). Mean filter and background subtraction was applied for clarity. The image shows a representative image from one of three independent experiments. See Figure S5 for additional data. **(b-d)** Infection time course of click labeled HIV-1* CA14^SiR^ compared to HIV-1*. Quantification of cells containing nuclear CA positive objects at the indicated times post infection for HIV-1* (black; CA(IF)), HIV-1* CA14^SiR^ (red; CA(IF)/CA(SiR)) and noninfected control (grey; CA(IF)). Mean values and SD from three independent experiments are shown (n>115 cells per timepoint). **(c)** Number of nuclear CA foci per cell determined for cells infected with HIV-1* (black) or HIV-1* CA14^SiR^ (red) at the indicated timepoints. n>120 cells were analyzed per sample. The median and interquartile lines are indicated in grey. **(d)** Localization of particles within the cell for HIV-1* (black) and HIV-1* CA14^SiR^ (red). The proportion of total particles per cell detected at the PM or in the cytoplasm (= PM/cytoplasm) at the nuclear envelope (=NE assoc.) or inside the nucleus was determined at the indicated time points. Data from two of three independent experiments are shown. Graphs show median of analyzed field of views in red (n>20 cells per condition) and interquartile lines.

Next, TZM-bl cells infected with HIV-1* or HIV-1*CA14^SiR^ were fixed and analyzed for the presence of click labeled subviral particles inside the nucleus at different time points after infection. Consistent with earlier results (Zurnic 2020, Burdick 2020, Müller 2021), we observed nuclear CA(IF) positive foci in HIV-1* infected cells as early as 6 h p.i. (Figure 4b, black), while such signals were absent in noninfected cells (Figure 4b, grey). Importantly, we detected SiR positive complexes in the nucleus of HIV-1*CA14^SiR^ infected cells, with the vast majority also positive for CA(IF) (Figure 4b, red). Nuclear entry appeared to be delayed for HIV-1*CA14^SiR^ compared to HIV-1* by 6-12 h. Nevertheless, samples infected with HIV-1*CA14^SiR^ displayed a comparable proportion of cells with detectable capsid-like objects in the nucleus as the HIV-1* infected control at 12 h p.i. (Figure 4b). The same was true for the number of nuclear objects detected per cell (Figure 4c); at 12 h p.i., HIV-1*CA14^SiR^ reached an average of 4.58 ± 4.12 nuclear particles per cell, similar to HIV-1* with 5.91 ± 4.11.

Delayed detection of subviral complexes in the nucleus may be due to slower uptake, slower trafficking towards the nuclear envelope, delayed passage through the NPC, or a combination thereof. In order to distinguish between these possibilities, we extended the time-resolved quantification to objects in close vicinity to the nuclear envelope (Figure 4d). This analysis revealed that the HIV-1*CA14^SiR^ derived subviral structures reached the nuclear envelope with similar kinetics as HIV-1* particles (Figure 4d, NE assoc.). A comparable average proportion of CA containing objects was detected at the nuclear envelope in both cases at 6 h p.i., while the numbers of nuclear capsids were lower for HIV-1*CA14^SiR^ at that time (Figure 4d, Nucleus). In contrast, the highest proportion of HIV-1*CA14^SiR^ nuclear objects with 4.20 ± 1.80% was detected at 18 h p.i., while HIV-1* reached similar levels at 6 h p.i. This implies that the mechanistic action of the capsid in nuclear import underlies tight margins with respect to its biophysical properties.

### Characterization of nuclear CA^SiR^ containing complexes

A long-standing question in the field of HIV-1 early replication is when and where genome uncoating takes place. The possibility to directly detect CA molecules clicked to a synthetic fluorophore enabled us to assess the amounts of CA associated with subviral complexes at different intracellular sites, without the influence of differential epitope accessibility or of a tag domain that potentially confers different properties to a subpopulation of CA molecules. However, comparing labeling intensities for nuclear, cytoplasmic, and extracellular particle-associated structures may be additionally confounded in diffraction-limited microscopy by the failure to resolve closely adjacent individual capsids. Clusters of nuclear capsids had indeed been observed by CLEM analyses in our previous study (Muller et al., 2021).

To determine whether nuclear cluster formation occurred under our conditions, we exploited the fact that the chemical dye conjugated to the capsid surface renders the modified virus suitable for super-resolution microscopy. With a lateral resolution of <50 nm, STED nanoscopy allows visual separation of closely adjacent CA objects. TZM-bl cells were infected with HIV-1*CA14^SiR^ at two different MOIs. MOI∼0.8 corresponded to the conditions generally used in our experiments; a 10-fold higher virus dose (MOI ∼8) was applied in a parallel experiment to test for enhancement of capsid clustering. At 18 h p.i., cells were fixed, immunostained against CA, and imaged using a STED system in confocal and STED mode (Figure 5). Nuclear CA(IF)/(SiR) double-positive objects were detected under both conditions (Figure 5a, arrowheads). While these objects appeared as individual punctae in diffraction-limited micrographs from the IF and SiR channels at both MOIs (Figure 5b, top and middle row), imaging of the SiR channel in STED mode revealed differences between individual punctae. Some diffraction-limited punctae in the nucleus represented individual capsid-like objects when imaged by STED (Figure 5b, left panel, bottom row). In contrast, other punctae were resolved into small clusters of 2-4 closely apposed CA-containing objects by super-resolution microscopy (Figure 5b, right panel, bottom row), consistent with observations made by electron tomography (Muller et al., 2021; Zila et al., 2021a). A quantitative analysis of cluster sizes (Figure 5c) revealed that the propensity for capsid clustering in the nucleus correlated with the amount of virus used for infection. Consistent with a normal distribution of infection events per cell at a given MOI, ∼50% of those cells positive for nuclear punctae displayed only a single STED resolved object at MOI 0.8, whereas all cells analyzed at MOI 8 comprised more than one STED resolved nuclear object, with up to 15 objects identified in one nucleus (Figure S6). Accordingly, at MOI∼0.8, the vast majority of punctae (∼88%) corresponded to individual capsid-like objects in the nucleus, and clusters of more than two objects were not observed. In contrast, the majority of cells analyzed comprising more than one nuclear object displayed clustered particles, and almost half of the nuclear punctae (∼43%) corresponded to clusters of 2-4 objects when cells were infected with MOI∼8. We conclude that nuclear capsid clustering is rarely observed at the MOI of 0.8 used throughout this study, and that nuclear clustering appears not to be a random event. The previously observed capsid clustering in distinct nuclear positions as well as larger clusters occurred preferentially at high MOI, where a higher proportion of cells contained multiple nuclear particles.

**Figure 5.**
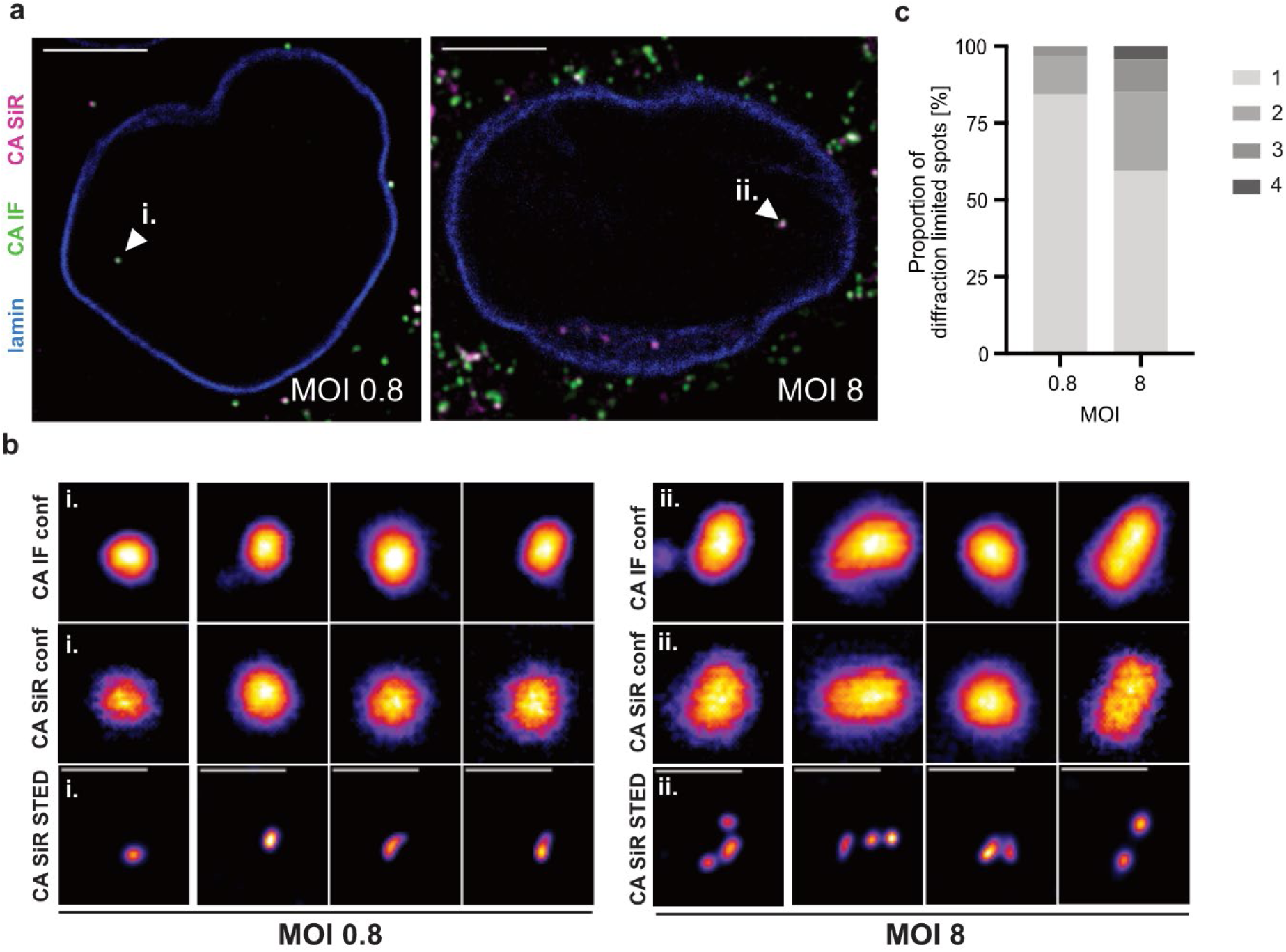
Dose dependent clustering of nuclear capsids in HeLa-derived cells. TZM-bl cells were infected with HIV-1* CA14^SiR^ at the indicated MOI, treated with 15 µM PF74 for 1 h before fixation at 18 h p.i., immunostained against CA (green) and lamin A/C (blue) and imaged using an Abberior STED microscope setup. Mean filter and background subtraction was applied to all images for clarity. **(a)** Micrographs of TZM-bl cells infected with MOI ∼0.8 (left) or MOI ∼8 (right). Arrowheads indicate nuclear CA(IF)/CA(SiR) positive objects shown enlarged in (b). Scale bars: 10 µm. **(b)** Representative images of nuclear CA containing objects from cells infected with a low MOI (MOI∼0.8, left panel) or a high MOI (MOI∼8, right panel). CA(IF) and CA(SiR) were imaged in confocal mode (top and middle row, respectively). CA(SiR) images were also recorded in STED mode (bottom row). The figure shows four representative foci each from one of two individual experiments. Mean filter and background subtraction were applied. Scale bars: 500 nm. **(c)** Diffraction-limited nuclear foci were analyzed by STED nanoscopy in cells infected with MOI∼0.8 (n = 32 foci) or MOI∼8 (n =47 foci) and classified by the number of individual capsids per focus.

We next proceeded to SiR fluorescence intensity measurements, comparing the signal intensity of extranuclear HIV-1 particles to that of subviral structures in the nucleus. Staining of the plasma membrane with mCLING ATTO488 before infection revealed that under our conditions most cell-associated particles in the cytosolic region represented virions present in endosomes, corresponding to a pre-fusion state of the virus (Figure S7a-c). To ensure that these extranuclear punctae represented single objects, cytoplasmic foci were analyzed in STED mode. We found that ∼95% (n=79) of analyzed punctae corresponded to an individual object, while only ∼5% (n=4) of these foci were resolved into two adjacent objects by nanoscopy (Figure S8).

As illustrated by the cartoon in Figure 6a, complete virions comprise on average ∼2,400 CA molecules, while only ∼1,200-1,500 of these are part of the mature fullerene capsid (Briggs 2003, Carlson 2008, Lanman 2004) that represents a post-fusion state. Assuming equal click labeling efficiency of CA14^CpK^ for molecules that are part of the mature lattice and those that remain free in the virion volume, the average SiR intensity of complete capsids is expected to correspond to ∼50-60% of the average intensity of complete virions from the same preparation. We infected TZM-bl cells at an MOI of 0.8 and quantified the SiR intensity of virions attached to the cell or in the cytosolic region (and thus mostly in endosomes) as well as nuclear punctae (Figure 6b). As a control, we compared SiR intensities of the small proportion of mCLING-negative (post-fusion) structures in the cytoplasmic region with that of mCLING positive (endosomal) objects. This analysis revealed an average SiR intensity of ∼ 66% for mCLING negative compared to mCLING positive objects (Figure S7d), indicating that we can reliably differentiate between complete virions and post-fusion complexes based on intensity of CA staining. The average SiR intensity of >6,000 cell-attached and (mostly) endosomal particles in the cytosolic region exhibited an average of 17,649 a.u.. In contrast, the SiR intensity of >100 nuclear subviral structures averaged 9,835 a.u. (Figure 6b), i.e., ∼ 56% of the average intensity of complete virions, similar to the predicted relative CA content of the mature capsid and similar to the intensity of mCLING-negative cytoplasmic objects. Based on these findings, we conclude that the CA(SiR) containing objects in the nuclei of these cells correspond approximately to the full CA complement of the mature capsid.

**Figure 6.**
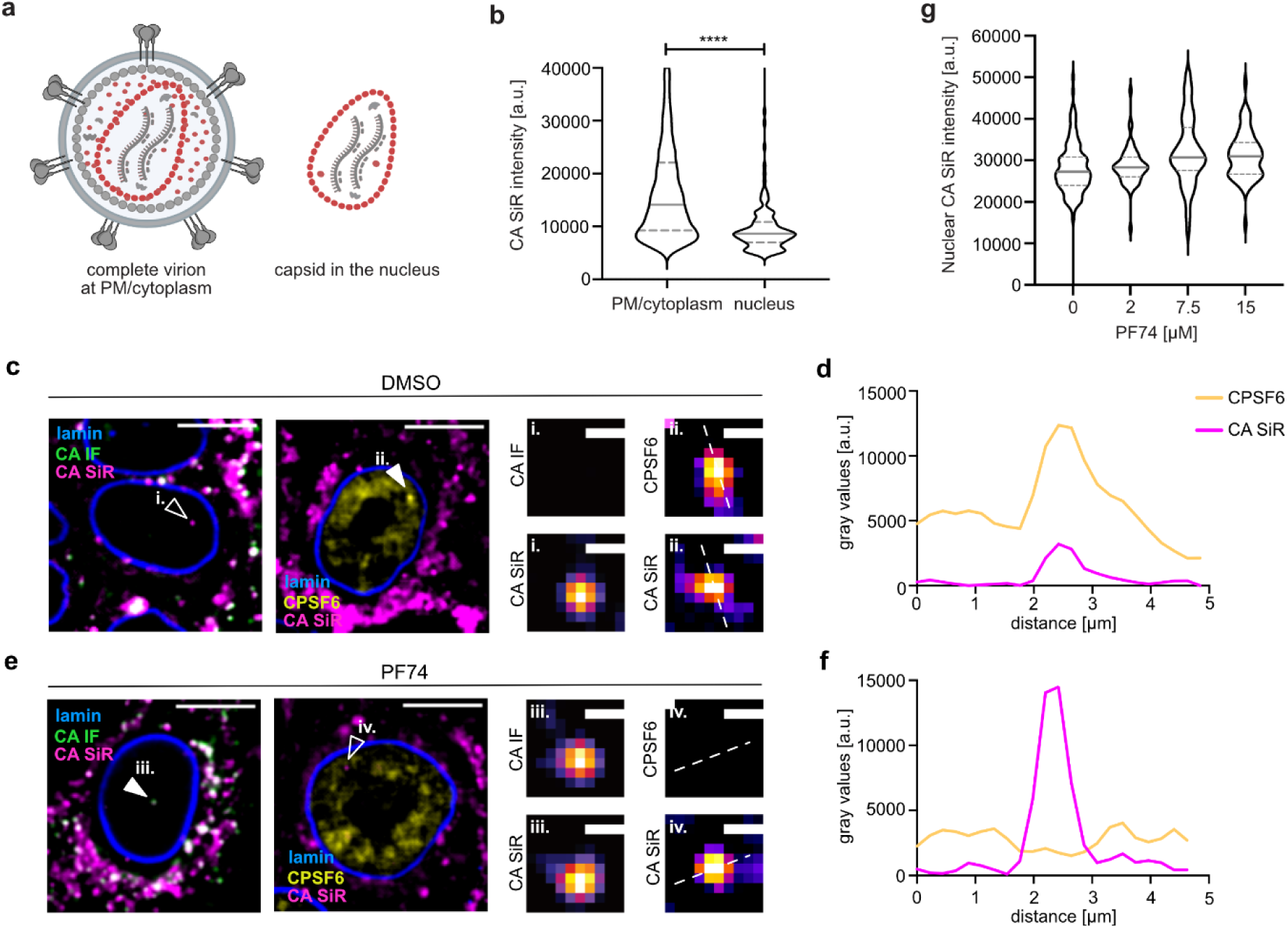
Largely intact capsids are detected in the nucleus of HeLa-derived cells. TZM-bl cells were infected with HIV-1* or HIV-1* CA14^SiR^ (MOI∼0.8), treated with 15 µM PF74 for 1 h before fixation at 18 h p.i. and imaged by SDCM. **(a)** Scheme of the relative CA content in complete virions (∼2,400 CA) on glass/plasma membrane or in endosomes in the cytosol. Post-fusion capsids contain only the CA molecules incorporated into the mature capsid lattice (∼1,500 CA). Image created with BioRender. **(b)** Quantification of CA(SiR) intensities associated with CA(IF) positive objects at the indicated localizations. Data from three independent experiments are shown. Cells from 7 fields of view were analyzed (nparticles= 6,441 PM/cytoplasm, 135 nucleus). Lines indicate median values (PM/cytoplasm: 17,649.22 ± 11,663.47; nucleus: 9,835.08 ± 5,708.14) and interquartile range. Significance was determined by two-tailed Student’s t-test (*** < 0.001). **(c-g)** TZM-bl cells were infected with HIV-1*CA14^SiR^, treated with DMSO **(c,d)** or 15 µM PF74 **(e,f)** for 1h before fixation at 18 h p.i.. Cells were immunostained against lamin A/C and HIV-1 CA or CPSF6 and subsequently imaged by SDCM. Scale bars: 10 µm (cells) and 1 µm (enlargements). **(d)** Quantification of intensity profile measured across the dashed line in **(c,ii.)** for CA(SiR) and CPSF6. **(f)** Quantification of intensity profile measured across the dashed line in **(e,iv.)** for CA (SiR) and CPSF6. **(g)** Quantification of CA(SiR) intensities of nuclear objects. TZM-bl cells were treated with DMSO (n=115) or 2 µM (n=58), 7.5 µM (n=110), 15 µM (n=69) PF74 for 1 h prior fixation at 17 h p.i.. Graphs show median values (DMSO: 2,7255; 2 µM: 2,8334.51; 7.5 µM: 3,0749.45; 15 µM: 3,0957) and interquartile range.

The small molecule inhibitor PF74 (Blair et al. 2010) binds to the HIV-1 capsid in a pocket overlapping the binding sites for the FG motifs of various nucleoporins and for the nuclear host protein CPSF6 (Bhattacharya et al., 2014; Matreyek et al., 2013; Price et al., 2012). This compound inhibits HIV-1 replication by multiple mechanisms, with effects on different replications steps reported based on PF74 concentration and time of addition (reviewed in (Thenin-Houssier and Valente, 2016)). Treatment with high concentrations of PF74 has been reported to destabilize the capsid (Burdick et al., 2020; Shi et al., 2011), but data obtained in recent studies argued against a PF74 induced loss of CA from nuclear complexes (Muller et al., 2021) and even showed an increased CA(IF) signal (Francis et al., 2020b). These findings are consistent with a stabilizing effect of the compound on the capsid lattice observed for isolated capsids after initial opening (Marquez et al., 2018; Rankovic et al., 2018), and for *in vitro* assembled CA-NC particles (Dostalkova et al., 2020). Since results obtained by immunodetection may be influenced by differential CA epitope exposure, we revisited this issue employing direct CA labeling. TZM-bl cells were infected with HIV-1*CA14^SiR^ particles for 17 h and then treated with 2-15 µM PF74 or DMSO for 1 h, followed by fixation, permeabilization, methanol extraction, and SDCM imaging. CA(SiR) punctae detected in the nucleus of DMSO control cells were positive for CPSF6 (Figure 6c ii). Intensity profile measurement revealed clear colocalization of both signals indicating that direct capsid labeling did not affect association with CPSF6 (Figure 6d). In accordance with earlier results (Francis et al., 2020b; Muller et al., 2021), CA(SiR) positive subviral complexes in nuclei of cells subjected to PF74 treatment lacked CPSF6 association (Figure 6e, and 6f). Their mean CA(SiR) intensity remained unaltered, however, indicating that the capsid remains largely stable in the presence of PF74 concentrations ranging from 2-15 µM (Figure 6g).

### Detection of directly labeled HIV-1 capsids in primary cells

To validate our results in a physiologically relevant cell type, primary human CD4^+^ T cells from healthy blood donors were infected, subjected to IF staining against CA, and imaged by SDCM at 24 h p.i. (Figure 7a and Figure S9). We readily detected nuclear subviral SiR positive structures in HIV-1*CA14^SiR^ infected cells, indicating that nuclear replication complexes retained CA also in these primary cells (Figure 7a). Consistent with prior observations made in T cell lines (Zila et al., 2019) the majority of SiR-positive objects were not associated with CA(IF) signals (9/11 particles; Figure 7a, left and Figure S9a) when fixation and immunostaining were performed under standard conditions. As outlined above, treatment with 15 µM PF74 for 1 h dissociates the large clusters of CPSF6 from nuclear subviral complexes. We observed that this in turn renders nuclear CA accessible for IF detection in T cells, presumably by exposure of CA epitopes upon CPSF6 displacement (Muller et al., 2021). Accordingly, brief PF74 treatment allowed for detection of CA(IF) signals co-localizing with nuclear CA(SiR) punctae (13/16; Figure 7a, right and Figure S9b). We conclude that the direct CA labeling strategy presented here overcomes technical artifacts that hamper IF analyses.

**Figure 7.**
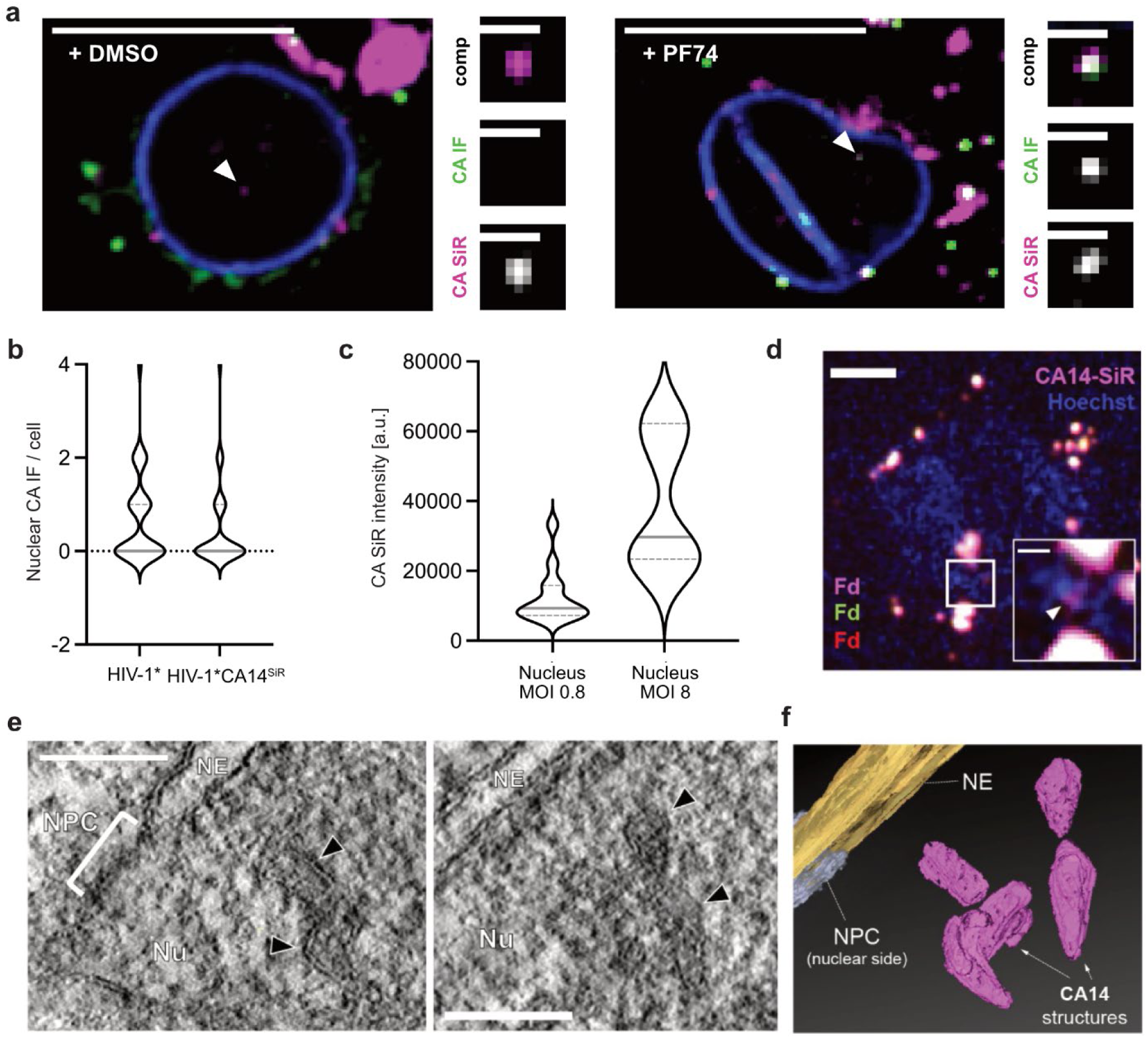
Largely complete click labeled capsid structures detected in the nucleus of primary CD4^+^ T cells and T cell line. **(a)** Activated CD4^+^ T cells were infected with HIV-1*CA14^SiR^ (MOI∼0.8), treated with DMSO/PF74 treatment for 1 h before fixation at 24 h p.i., and extracted with methanol. Samples were immunostained against CA (green) and lamin A/C (blue). Images show a single z slice through the cell. Enlargements show the particle marked by the arrowhead. Scale bars: 10 µm (overview) and 1 µm (enlargement). **(b)** Data analyzed from the experiment outlined in (a). The graph shows the number of CA positive foci per nucleus in cells infected with HIV-1* (n=35 cells, mean=0.85) or HIV-1*CA14^SiR^ (n= 73 cells, mean=0.51). Pooled data from 6 different blood donors are shown. Grey lines show median and interquartile lines. **(c)** CA(SiR) intensities of nuclear objects in infected and activated CD4^+^ T cells at an MOI∼0.8 (n=13; mean=12,485 ± 7,445 a.u.) and an MOI ∼8 (n=7; mean=39,502 ± 18,025 a.u.). MOI was determined in TZM-bl cells. Grey lines show median and interquartile lines. **(d-f)** Nuclear cone-shaped capsids detected by CLEM-ET. SupT1 cells were treated with 1 µM aphidicolin (APC) for 16 h to prevent cell division, before infection with HIV-1*CA14^SiR^ virions (2.3 µU RT/cell, corresponds to an MOI∼0.4 determined in TZM-bl cells). At 24 h p.i., cells were cryo-immobilized by high-pressure freezing, freeze substituted, and further processed for CLEM and ET as described in materials and methods. **(d)** SDCM image of a 250-nm thick resin section of the cell infected with HIV-1*CA14^SiR^ virions (magenta), post-stained with Hoechst (blue) and decorated with multi-fluorescent fiducials (Fd) for correlation. The arrowhead in the enlargement of the boxed region indicates a CA(SiR) signal within the Hoechst-stained nuclear region. Scale bars: 1 µm (overview) and 200 nm (enlargement). **(e)** Computational slices through tomographic reconstructions at the correlated region boxed in (d) with views highlighting the presence of clustered capsid-reminiscent structures (black arrowheads) in the nuclear region. Nu, nucleus; NPC, nuclear pore complex; NE, nuclear envelope. Scale bar: 100 nm. **(f)** Segmented and isosurface rendered structure of the cones detected in (e). Magenta: capsid, yellow: NE, cyan: NPC (cryo-EM map of NPC: EMD-11967 (Zila et al., 2021a)). See also supplementary movie 1.

Further quantitative analyses using primary CD4^+^ T cells prepared from six blood donors revealed similar numbers of nuclear capsid structures in cells infected with HIV-1*CA14^SiR^ compared to cells infected with HIV-1* at 24 h p.i. (Figure 7b). SiR intensity measurements were only performed for intranuclear objects in this case, since high background due to SiR accumulation in the narrow cytoplasm of T cells precluded reliable analysis of individual particles in the extranuclear region. Quantitation of SiR intensities of nuclear punctae in cells infected with a MOI∼0.8 yielded similar average intensities as measured in TZM-bl cells (mean=12,485 a.u.), indicating the presence of a complete or nearly complete mature capsid in the nuclear complexes in primary T cells as well (Figure 7c). Cells infected with a MOI of ∼8 displayed higher CA(SiR) intensities of diffraction-limited nuclear objects (mean=39,502 a.u.), suggesting intranuclear clustering of capsids, as observed in TZM-bl cells (Figure 5).

Our findings from CA(SiR) intensity measurements argued for the presence of a full capsid complement at subviral structures in the nucleus. These data strengthen conclusions from several recent studies suggesting that the mature capsid lattice may be completely or largely intact on nuclear subviral objects (Burdick et al., 2020; Muller et al., 2021; Zila et al., 2021a). However, fluorescence signals do not yield information on the architecture of nuclear CA14^SiR^ containing objects. Therefore, we complemented our analyses by performing CLEM of infected SupT1 T cells. In order to maximize the number of nuclear objects, infection was synchronized by the attachment of particles to the cells for 3 h at a low temperature (16°C) to prevent particle uptake by membrane fusion or endocytosis (Melikyan et al., 2000; Weigel and Oka, 1981). Virus entry was initiated by temperature shift to 37°C. At 24 h post temperature shift, specimens were prepared by high-pressure freezing and freeze substitution. 250 nm thick resin sections were subjected to SDCM in order to localize CA(SiR) containing structures, followed by correlative electron tomography (CLEM-ET) analysis. CA(SiR) positive objects could be identified by SDCM in the sections (Figure 7d), demonstrating that the brightness of signals derived from direct CA(SiR) labeling is sufficient for CLEM detection of cytosolic and nuclear (sub)viral structures. ROIs were defined based on the SiR signals and subjected to correlative ET analysis. Figure 7e shows an exemplary tomogram obtained from a ROI located within the nucleus. It reveals several closely apposed electron-dense structures at the position of the SiR label, whose shape and dimension match those of intact or largely intact mature HIV-1 capsids (Figure 7f and supplementary movie 1). Such structures were recently identified in nuclei of infected cells by CLEM using fluorescently labeled HIV-1 IN as an indirect marker for subviral structures (Muller et al., 2021; Zila et al., 2021a) and were interpreted as capsid shells solely based on their morphology. Here, we demonstrated that these capsid-resembling structures co-localize with nuclear foci comprising a high number of click labeled CA molecules. We thereby provide direct evidence that the cone-shaped objects are complete or largely complete HIV-1 capsids that have entered the nucleus presumably through the NPC of infected and cell cycle arrested cells.

## Discussion

Here, we present a direct labeling approach for HIV-1 CA that yields infectious and morphologically mature viral particles. The minimally invasive GCE/click labeling approach used here represents an ideal strategy for the versatile labeling of genetically fragile viral capsid proteins in principle, but its potential for virus imaging has not been exploited so far. The combination of GCE and subsequent functionalization of a viral capsid protein by click chemistry has previously only been applied to the non-enveloped adeno-associated virus (AAV) (e.g., (Kelemen et al., 2018; Zhang et al., 2018). However, the capsid of AAV, unlike HIV-1 CA, can also tolerate peptide insertions and larger modifications (Borner et al., 2020; Chandran et al., 2017; Feiner et al., 2019; Varadi et al., 2012). Here, we demonstrate that GCE in conjunction with click labeling can also be applied to an enveloped virus with a highly multifunctional and genetically fragile capsid protein that needs to form a closed fullerene lattice to be infectious. We found that HIV-1 CA tolerates chemical modification of the exposed amino acid residue A14. This position is located close to the central, dynamic pore in the CA hexamer lined by positively charged R18 and K25 rings (Jacques et al., 2016). Replacement of A14 and E45 by cysteine residues for disulfide cross-linking CA subunits was previously shown to allow CA hexamer assembly and was used to study the capsid hexamer structure (Pornillos et al., 2009). This mutant retained binding to host cell proteins Nup153 and CPSF6, as well as to dNTPs and PF74 (Jacques et al., 2016; Price et al., 2014), consistent with functional replacement of A14 by a non-natural CpK residue in the current study.

All previously described genetic tagging strategies for HIV-1 CA (Burdick et al., 2020; Campbell et al., 2008; Pereira et al., 2011; Zurnic Bonisch et al., 2020) required complementation with a molar excess of wt protein or virus. Since the mature HIV-1 capsid is assembled from approximately half of the ∼2,500 CA molecules packaged in the virion (Briggs et al., 2004; Lanman et al., 2004), it cannot be ascertained in this case whether the subset of genetically tagged CA molecules is an integral part of the mature capsid lattice. Conceivably, tagged or modified CA molecules may be excluded from the mature capsid shell or may be irregularly inserted. Signal intensity changes of CA fusion proteins in early HIV-1 replication may thus also depend on the relative incorporation of these proteins into the mature capsid lattice. In contrast, we found that the strategy described here allowed genetic labeling of HIV-1 CA in the proviral context and retaining almost full infectivity in the absence of wild-type complementation.

The detection of a label covalently attached to CA is independent of cellular context, sample treatment, or exposure of CA epitopes. Thereby, the method overcomes limitations of IF detection that had previously resulted in different conclusions regarding the presence of CA on subviral complexes. The use of synthetic dye molecules also renders the labeling strategy compatible with a wide range of fluorescence imaging approaches, including live-cell microscopy, correlative imaging and super-resolution fluorescence microscopy techniques (Wang et al., 2019).

Our approach allowed for direct, quantitative analysis of CA-containing objects and CA amounts associated with viral complexes in microscopic images of infected cells. While time-lapse experiments showed some delay in nuclear import kinetics for labeled capsid-like objects, the infectivity of highly labeled preparations was reduced by only twofold, and the number of nuclear objects reached was similar to that detected in cells infected with wt virus. Thus, site-specific introduction of a synthetic fluorophore can be compatible with capsid functionality in HIV-1 post-entry processes. The delay in nuclear accumulation appears to be mainly caused by slower trafficking through the NPC, possibly due to the additional mass of the ncAA and label, given that the unmodified HIV-1 capsid already approaches the size limit even of dilated NPCs (Zila et al., 2021a). Another, not mutually exclusive, possibility is that ncAA incorporation and/or the attached SiR dye affects pliability of the capsid structure that might be required for efficient and fast passage through the NPC.

CA amounts approximately corresponding to the full complement of a mature capsid were found to be associated with cytoplasmic post-fusion objects and with subviral complexes in nuclei of a HeLa-derived cell line and primary human CD4^+^ T cells, also upon inhibition of cell division by aphidicolin treatment. By applying correlative imaging, we provide direct evidence that nuclear complexes positive for directly labeled CA indeed represent HIV-1 capsids or capsid-like remnants. Taken together, these results argue against (partial) capsid uncoating prior to entering the nucleoplasm, as had been concluded earlier based on low or lacking CA IF signals associated with nuclear subviral complexes in certain cell types (e.g. (Burdick et al., 2017; Hulme et al., 2015; Peng et al., 2014; Zila et al., 2019), or based on the loss of the fluorescently labeled capsid binding protein CypA at the nuclear envelope (Francis et al., 2020a; Francis et al., 2016; Francis and Melikyan, 2018). The apparent discrepancy between these previous IF results and data from direct CA quantification may be explained by differential accessibility of CA epitopes under different IF conditions. The indirect label CypA, on the other hand, might be displaced from capsids at the nuclear pore, possibly by competition between fluorescent CypA and the outer NPC protein Nup358, which also carries a binding site for the CypA binding loop of CA (Schaller et al., 2011). Our data suggest nuclear capsid uncoating in a model cell line, as well as in primary T cells, in agreement with recent findings from us and others, which indicated that the nuclear pore channel is wider than previously assumed, allowing HIV-1 capsids to pass the intact NPC (Zila et al., 2021a), and that HIV-1 uncoating occurs after nuclear import (Burdick et al., 2020; Dharan et al., 2020; Li et al., 2021; Muller et al., 2022; Selyutina et al., 2020), apparently by separation of the viral genome from a broken capsid remnant (Muller et al., 2021).

Small clusters of CA positive objects were detected by STED nanoscopy in nuclei of TZM-bl cells and T cells, consistent with the reported detection of nuclear clusters containing multiple HIV-1 replication complexes (Francis et al., 2020b), multiple viral genomes (Rensen et al., 2021), or even several intact or partly intact capsid-like structures (Muller et al., 2021) in various cell types. Our analyses revealed that the observed clustering is dependent on the amount of virus used for infection. Most nuclear signals represented single capsids at a lower MOI, whereas frequent clustering was observed at high MOI. This observation suggests that capsids enter the nucleus individually, but traffic via a limited number of routes and accumulate at defined sites of uncoating. This raises the question whether HIV-1 capsids use a ‘specialized’ subset of nuclear pores for nuclear entry; the answer would not only be relevant in the context of HIV-1 replication, but also with respect to an understanding of the nuclear import process. Intracellular Nup levels and presumably NPC composition have been reported to influence HIV-1 replication (Kane et al., 2018), but compositional and structural variability of NPCs between different cell types, or within an individual cell, is incompletely understood (reviewed in (Knockenhauer and Schwartz, 2016). The route, mechanism and functional consequences of intranuclear trafficking of HIV-1 complexes also warrant further analysis. Growing evidence from recent studies suggests that incoming viral replication complexes accumulate at nuclear speckles in a CA and CPSF6-dependent manner, and that reverse transcription may only be completed near the site of integration (Burdick et al., 2020; Francis et al., 2020a; Rensen et al., 2021; Selyutina et al., 2020). Combining the direct CA labeling described here with the recently developed fluorescence detection of the reverse transcribed genome (Blanco-Rodriguez et al., 2020; Muller et al., 2021) will provide us with the possibility to study the uncoating process in more detail using a combination of confocal imaging, nanoscopy and correlative imaging.

The direct labeling approach also allowed us to investigate the effect of the capsid inhibitor PF74 (Blair et al., 2010), whose detailed mode of action is still under investigation. We found that displacement of CPSF6 from nuclear subviral structures was not accompanied by a loss of CA signal. This finding disagrees with the recently reported rapid CA dissociation from nuclear complexes upon PF74 addition, that was based on imaging of HIV-1 particles containing eGFP-CA complemented by a molar excess of wt CA (Burdick et al., 2020). The apparent discrepancy may suggest that the subset of eGFP-tagged CA molecules is not an integral part of the mature capsid lattice, resulting in premature loss of the labeled molecules. Our findings are in line with the observation of other recent studies where PF74 treatment did not lead to a loss of the CA IF signal on nuclear complexes but rather enhanced immunostaining efficiency (Francis et al., 2020b; Muller et al., 2021) and with *in vitro* findings that indicated breach of lattice integrity by PF74, but stabilization of the remaining lattice (Marquez et al., 2019; Rankovic et al., 2018).

In conclusion, direct click labeling of HIV-1 CA is a versatile approach that substantially expands the possibilities to study the early events in HIV-1 replication with high temporal and/or spatial resolution using advanced fluorescence microscopy methods. Application for HIV-1 CA demonstrated its usefulness for genetically fragile structural proteins and provided direct proof that the capsid stays largely intact upon passage of the subviral complex into the nucleus. Furthermore, it directly identified nuclear capsid-like structures that morphologically resembled the virion capsid by CLEM-ET. The variety of synthetic fluorophores which can be combined with GCE and click labeling enables rapid adaptation of the approach to microscopic single-molecule techniques such as STORM (Rust et al., 2006) and MINFLUX (Balzarotti et al., 2017). It can also be combined with labeling of proteins of the viral replication complex and of the viral genome, which is a subject of ongoing studies. Beyond fluorescence imaging, CA modification by GCE offers possibilities for site-specific incorporation of other functionalized ncAAs, e.g., benzoyl-phenylalanine (Coin, 2018) for photo-crosslinking, together with pulldown experiments for the detection of capsid-host protein interactions. The fact that the combination of GCE and click chemistry could successfully be applied to a notoriously genetically fragile capsid protein of an enveloped virus opens the perspective that this strategy may also advance and expand fluorescence labeling of a broad range of other viruses.

## Acknowledgements

We acknowledge microscopy support from the Infectious Diseases Imaging Platform (IDIP) at the Center for Integrative Infectious Disease Research, Heidelberg. We thank Eyal Arbely (The National Institute for Biotechnology in the Negev, Beer-Sheva, Israel) for providing pEA168. pNESPylRS-eRF1dn-tRNA was kindly provided by Volkan Sakin and Anna-Lena Schäfer (University Hospital Heidelberg, Heidelberg, Germany).

This work was funded in part by the Deutsche Forschungsgemeinschaft (DFG; German Research Foundation), Projektnummer 240245660, SFB 1129 project 6 (B.M.) and project 5 (H.G.K.) and from DZIF (to H.G.K. and V.L.).

## Materials and Methods

### Plasmids

Plasmids were cloned using standard molecular biology techniques and verified by commercial Sanger sequencing (Eurofins Genomics). PCR was performed using Q5 High-Fidelity DNA Polymerase (New England Biolabs) or Phusion DNA Polymerase (New England Biolabs) according to the manufacturer’s instructions using primers purchased from Eurofins Genomics. Plasmid amplification was carried out in *E. coli* Stbl2 (Thermo Fisher Scientific) cells.

HIV-1 plasmids were based on the proviral plasmid pNLC4-3 (Bohne and Krausslich, 2004) that expresses the authentic genomic RNA from HIV-1_NL4-3_ (Adachi et al., 1986) under the control of the cytomegalovirus promoter. To avoid unwanted ncAA incorporation into the virion component Vpr, the amber stop codon of the *vpr* ORF of pNLC4-3 was mutated into an opal stop codon (TGA) via site-directed mutagenesis. See primer list for sequences of primers used. PCR1 (primers Vpr_TGA_ a and Vpr_TGA_ b) and PCR2 (primers Vpr_TGA_ c and Vpr_TGA_ d) were performed in parallel to generate two overlapping single stranded PCR products. Using a combination of both products of these reactions as new templates, PCR3 with primers Vpr_TGA_ a and Vpr_TGA_ d resulted in PCR fragments comprising the respective mutation. These fragments were cloned into pNLC4-3 using unique PflMI/NheI restriction sites, resulting in pNLC4-3*. Virus produced from this plasmid was termed HIV-1*.

To allow for site-specific GCE, the codon for amino acid A14 of CA was mutated into TAG via overlap PCR. PCR1 (primers CA14_BssHII_ fwd 1, CA14_TAG_ rev 1) and PCR2 (primers CA14_TAG_ fwd 2, CA14_ApaI_ rev 2) were performed in parallel to generate two overlapping single-stranded PCR products. PCR3 with primers CA14_BssHII_ fwd 1 and CA14_ApaI_ rev 2 result in the PCR fragment comprising the mutation, which was subcloned into pNLC4-3* using unique BssHII/ApaI restriction sites, resulting in pNLC4-3*CA14^TAG^ (HIV-1*CA14^TAG^).

Plasmid pNESPylRS-eRF1dn-tRNA (Schifferdecker, Sakin et al., in preparation) is based on pEA168 (Cohen and Arbely, 2016), kindly provided by Eyal Arbely, Ben-Gurion University of the Negev, Israel), a eukaryotic vector that comprises expression cassettes for two proteins and four tRNA molecules. The coding sequence for a modified pyrrolysine tRNA synthetase was cloned from plasmid tRNA^Pyl^/NESPylRS^AF^ (Nikic et al., 2016) into the CMV promoter driven expression cassette of pEA168 using HindIII/XbaI restriction sites, resulting in plasmid pEA168-CMV-aaRS-4xU6tRNA. A PCR fragment encoding a dominant version of the eukaryotic release factor 1 (eRF1(E55D)) amplified from plasmid peRF1-E55D (Schmied et al., 2014) was subsequently inserted into an expression cassette driven by the EF1 promotor into pEA168-CMV-aaRS-4xU6tRNA using KpnI/MluI restriction sites, yielding pNESPylRS-eRF1dn-tRNA.

**Table 1.**
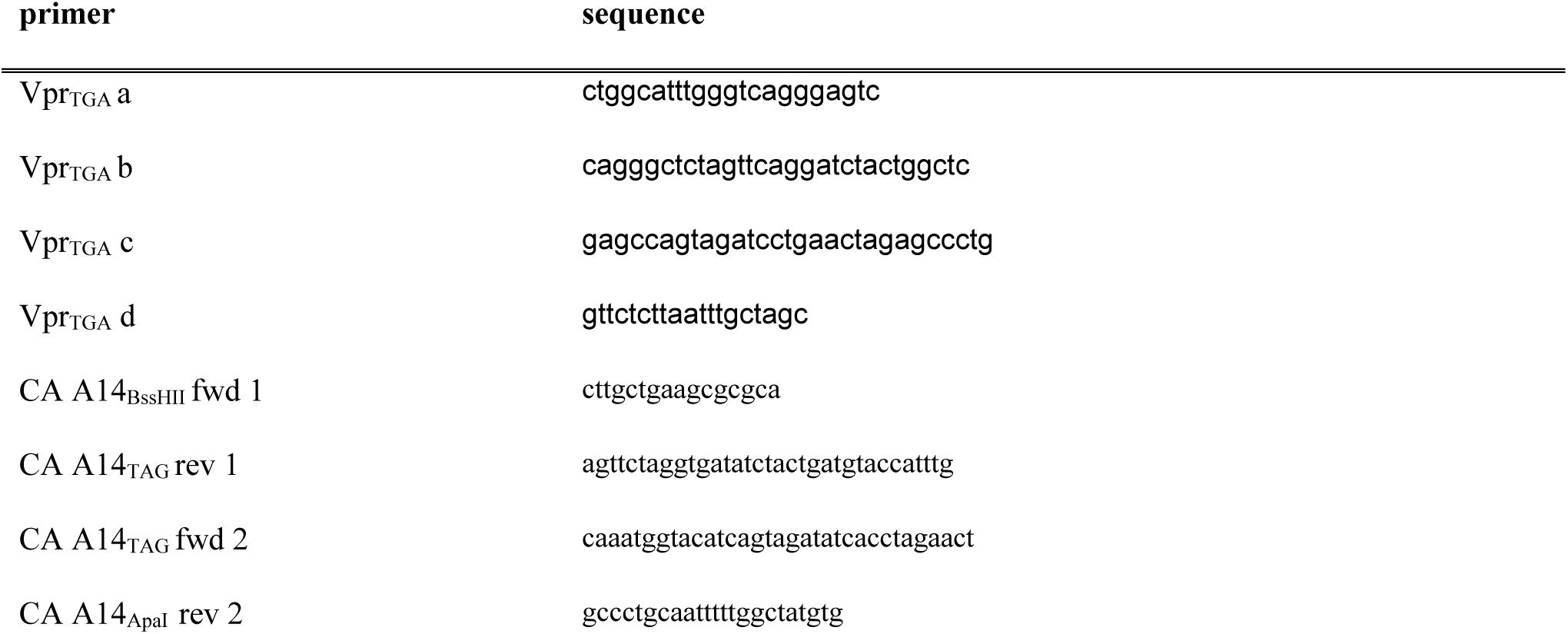
Primers

### Cell culture

HEK293T (Pear et al., 1993) and HeLa TZM-bl indicator cells (Wei et al., 2002) were maintained in Dulbecco’s Modified Eagle’s medium (Thermo Fisher Scientific) supplemented with 100 U/ml penicillin, 100 μg/ml streptomycin (PAN Biotech) and 10% fetal calf serum (FCS; Sigma Aldrich). Both cell lines were regularly monitored for mycoplasma contamination using the MycoAlert mycoplasma detection kit (Lonza Rockland). Primary CD4+ T cells were cultured in RPMI 1640 containing L-glutamine supplemented with 100 U/ml penicillin, 100 μg/ml streptomycin (PAN Biotech), 10% heat-inactivated FCS, and 5% human AB serum (Sigma Aldrich).

### Isolation of primary cells

Primary human CD4+ T cells were isolated from buffy coats obtained from healthy and anonymous blood donors at the Heidelberg University Hospital Blood Bank following the regulations of the local ethics committee. CD4+ T cells were isolated using EasySep^TM^ Direct Human T Cell Isolation Kit (Stemcell technologies) according to the manufacturer’s instructions and activated by incubation in the presence of 100 U/ml IL-2 (Sigma Aldrich) and T Cell TransAct™ human (Miltenyi Biotec) for 72 h.

### Virus particle production

HEK293T cells were seeded in T175 tissue culture flasks the day before (∼15 Mio. cells) or a 6-well plate (4x10^5^ cells/well) and transfected using calcium phosphate precipitation according to standard procedures (∼80% confluency). Cells were co-transfected with 50 µg/flask or 3 µg/well total DNA of pNLC4-3* (HIV-1*) or pNLC4-3*CA14^TAG^ (HIV-1*CA14^TAG^) and plasmid pNESPylRS-eRF1dn-tRNA in a molar ratio of 2.22:1. At 6 h p.t., medium was removed, and fresh complete DMEM containing a final concentration of 500 µM CpK (SiChem; stock solution of 100 mM was pre-diluted 1:4 in 1M HEPES shortly before use), and 100 µM ascorbic acid (Sigma Aldrich; stock solution 10 mM) was added. At 48 h p.t., the tissue culture supernatant was harvested and filtered through 0.45 µm nitrocellulose filters. For labeling the CA protein, 250 nM Tetrazine-SiR (Spirochrome; stock solution 1 mM) was added to the filtered supernatant, and samples were incubated at 37°C for 30 min. Particles were then concentrated by ultracentrifugation through a 20% (w/v) sucrose cushion at 28,000 rpm using a Beckman TLA-100 fixed angle-rotor (Beckman Coulter) for 90 min or for the 6-well format at 44,000 rpm using a Beckman TLA-55 fixed angle-rotor (Beckman Coulter) for 45 min at 4°C. Pellets were gently resuspended in phosphate-buffered saline (PBS) containing 10% FCS and 10 mM HEPES (pH 7.5) and stored in 5 µl aliquots at -80°C.

### Immunoblotting and In-gel fluorescence

Virus samples were mixed 1:10 with SDS sample buffer (150 mM Tris HCl, pH 6.8, 6% (w/v) SDS, 30% Glycerin, 0.06% bromophenol blue, 20% β -Mercaptoethanol) and boiled at 95°C for 15 min. 10 µl HIV-1* and 40 µl HIV-1*CA^SiR^ lysates were subjected to SDS-PAGE (15 %; acrylamide:bisacrylamide 200:1). Cell lysates were generated from transfected HEK293T cells. At 40 h p.t., cells were washed with PBS, trypsinized and resuspended in PBS. 1 ml of cell suspension was mixed with 300 µl SDS sample buffer and boiled at 95°C for 15 min. 10 µl cell lysate was subjected to SDS-PAGE. Proteins were transferred to a nitrocellulose membrane (Millipore) by semi-dry blotting for 1 h at 0.8 mA/cm^2^. Antigens were stained with the indicated antisera in PBS/0.5% bovine serum albumin (BSA) (sheep αCA, polyclonal 1:5,000 (in-house); rabbit αMA, polyclonal 1:1,000 (in-house); rabbit αRT, polyclonal, 1:1,000 (in-house), followed by staining with corresponding secondary antibodies IRDye^TM^ in PBS / 0.5% BSA (anti-sheep 680CW (1:10,000); Rockland, or anti-rabbit 800CW (1:10,000); Li-COR Biosciences). Detection was performed using a Li-COR Odyssey CLx infrared scanner (Li-COR Biosciences) according to manufacturer’s instructions. CA quantification was performed with ImageStudio LITE software (Li-COR Biosciences) via intensity measurements of CA bands and a serial dilution of recombinant purified CA standard (2.5 ng/µl; in-house) on the same membrane. For in-gel fluorescence, the acrylamide gels were directly scanned using a Li-COR Odyssey CLx infrared scanner (Li-COR Biosciences) set at an emission wavelength of 700 nm.

### Infectivity assays

Virus amounts were quantified via SYBR Green based Product Enhanced Reverse Transcription assay (SG-PERT; (Pizzato et al., 2009)). To determine relative infectivity of HIV-1* particles in a luciferase assay, 5,000 TZM-bl cells were seeded in 90 µl DMEM in a 96-well plate. On the following day, infection was performed 10 µl of serial 9-fold dilution of supernatant from virus producing HEK293T cells transfected with pNLC4-3 or pNLC4-3*. At 48 h p.i., cells were lysed by adding 100 µl Steady-Glo luciferase reagent (Promega). After 10 min, 80 µl of the lysed cell solution was transferred into a white 96-well plate and an Infinite 200 PRO plate reader (TECAN) was used to measure luminescence. Uninfected cells were used as a negative control.

To determine the effect of incorporating CpK and Tet-SiR labeling on virus infectivity, HIV-1* and HIV-1*CA14^SiR^ viral particles (normalized by RT activity) were titrated on TZM-bl cells seeded in 15-well ibidi µ-Slide angiogenesis dishes. At 6 h p.i. 50 µM T-20 (Enfuvirtide; Roche; stock solution 20 mM) was added to prevent second-round infection. Infection rates were scored at 48 h p.i.. For this, cells were fixed in 4% paraformaldehyde (PFA; Electron Microscopy Sciences; stock solution 16%) for 15 min, followed by 20 min incubation in PBS/0.5% (v/v) Triton X-100 at room temperature. Immunostaining was performed using an in-house polyclonal rabbit antiserum raised against recombinant HIV-1 MA (1:1,000; in-house) in PBS/0.5% BSA) 1 h at room temperature. Secondary antibody Alexa Fluor 488 donkey anti-rabbit (1:1,000; Thermo Fisher Scientific) in PBS/0.5% BSA was added for 45 min at room temperature. Samples were imaged by SDCM. The mean intensity of the 488 channel (MA IF) was quantified in the non-infected samples imaged in parallel and subtracted as background in each image. The proportion of IF-positive cells was counted in 12 randomly selected fields of view using Fiji (Schindelin et al., 2012). To determine the infectivity of virus particle preparations, the number of infected cells per well was calculated by multiplying the percentage of infected cells detected with the number of cells per well (double of seeded cell number the day before). Division by the volume of virus suspension used for infection yielded the number of infectious units (IU) / ml.

### Fixation and immunofluorescence staining of infected cells

3.33 × 10^3^ TZM-bl cells were seeded into 15-well µ-Slides angiogenesis dishes (ibidi; cat. 81507) the day before infection. Infection at 37°C was performed with an MOI ∼0.8 for 6,9,12 or 18 h. Subsequently, cells were incubated for 1 h with 15 µM PF74 (Sigma Aldrich; stock solution 10 mM in DMSO) in DMEM to allow for efficient detection of nuclear CA by IF (Muller et al., 2021). Samples were washed with PBS, fixed in 4% PFA for 15 min and permeabilized with PBS/0.5% (v/v) Triton-X100 for 20 min, and washed again with PBS. Cells were extracted using ice-cold 100% methanol for 10 min. Afterward, samples were blocked with PBS/2.5% BSA for 15 min, followed by incubation with primary antibodies (rabbit αCA, polyclonal 1:1000 (in-house); rabbit αCPSF6, polyclonal 1: (Atlas Antibodies, cat. #HPA039973); mouse αhLamin A/C, monoclonal 1:100 (Santa Cruz Biotech., cat. #sc-7292); mouse αhLaminB1, monoclonal 1:100 (Santa Cruz Biotech., cat. #sc-365962)) in PBS/0.5% BSA for 1 h at room temperature. After washing three times with PBS, secondary antibodies (Alexa Fluor 405, 488, 568, goat/donkey, polyclonal 1:1,000 (Thermo Fisher Scientific)) diluted in PBS/0.5% BSA were added for 45 min at room temperature. Samples were washed and stored in PBS at 4°C. For infection of primary CD4^+^ T cells, 20,000 cells were infected with HIV-1* or HIV-1*CA14^SiR^ in a 96-well v-bottom microplate (Greiner Bio-one, cat. #650161) in a volume of 40 µl RPMI and transferred at 22 h p.i. onto a PEI-coated 15-well µ-Slide angiogenesis dishes (ibidi). Cells were allowed to adhere for 1 h at 37°C, and PF74 diluted in fresh growth medium was added to a final concentration of 15 µM. Extraction, fixation, and immunostaining were performed after 1 h at 37°C as described above. For the detection of endosome-associated particles, 2 µM mCLING ATTO488 (Synaptic Systems; stock 50 µM) was added to TZM-bl cells seeded in 15-well µ-Slides Angiogenesis and incubated at 16°C for 30 min. Subsequently, the fluorescent probe was removed, HIV-1*CA14^SiR^ particles were added in fresh growth medium, and cells were incubated for an additional 3 h at 37°C (MOI∼0.8). Cells were fixed for 90 min at room temperature in 4% PFA and 0.2% glutaraldehyde to ensure retention of mCLING at cellular membranes. Nuclei were stained with 5 µg/ml Hoechst (Merck) in PBS for 30 min.

### Cell viability assay

To test the effect of mCLING ATTO488 (Synaptic Systems) staining on cell viability, TZM-bl cells were seeded into a 96-well plate (5x10^3^ cells/well; flat bottom Greiner Bio-one) the day before and incubated in medium supplemented with the indicated concentration of mCLING ATTO488 for 30 min at 16°C. After staining, cells were trypsinized, stained with Trypan blue using standard procedures and analyzed with a TC20^TM^ Automated Cell Counter (BioRad).

### Labeling efficiency of immobilized particles

15-well µ-Slide angiogenesis dishes (ibidi) were coated with 30µl/well polyethyleneimine (PEI; 1mg/ml) for 30 min at room temperature and washed with PBS. Pre-labeled HIV-1* and HIV-1*CA14^SiR^ particles were incubated in PBS on PEI-coated microscopy slides for 1 h at 37°C. Subsequently, samples were washed with PBS, fixed in 4% PFA for 15 min and permeabilized with PBS/0.05% (v/v) Triton X-100 for 20 min at room temperature. Immobilized particles were blocked with PBS/2.5% BSA for 15 min and polyclonal rabbit antiserum raised against recombinant HIV-1 CA protein (in-house) was added (1:1,000 in PBS/0.5% BSA for 1 h at room temperature). After washing three times with PBS, secondary antibody Alexa Fluor 488 donkey anti-rabbit (Thermo Fisher Scientific) 1:1,000 in PBS/0.5% BSA was added for 45 min at room temperature. Samples were washed and stored in PBS at 4°C.

### Confocal microscopy (SDCM)

Multichannel z-series with a z-spacing of 200 nm, spanning the whole cell volume (3D), were acquired using a PerkinElmer Ultra VIEW VoX 3D spinning disk confocal microscope (SDCM; Perkin Elmer). A 60x oil immersion objective (numeric aperture [NA] 1.49; Perkin Elmer) was used for imaging of TZM-bl cells or 100x oil immersion objective ([NA] 1.49; Perkin Elmer) for primary CD4+ T cells and immobilized particles. Images were recorded in the 405-, 488-, 561-, and 640 nm channels.

### STED microscopy

STED nanoscopy was performed using a λ = 775 nm STED system (Abberior Instruments GmbH) equipped with a 100x oil immersion objective (NA 1.4; Olympus UPlanSApo). STED images were acquired using the 640 nm excitation laser lines while the 488 and 590 laser line was acquired in confocal mode only. Nominal STED laser power was set to 20% of the maximal power (1250 mW) with pixel dwell time of 10 µs and 15 nm pixel size. STED images were deconvolved using the software Imspector (Abberior Instruments GmbH) and Huygens Professional Deconvolution (Scientific Volume Imaging).

### Electron microscopy

HEK293T cells (4×10^5^) were seeded in a glass coverslip-bottom petri dish (MatTek, MA, USA), cultured for 16 h at 37°C and then co-transfected with pNLC4-3*CA14^TAG^ and pNESPylRS-eRF1dn-tRNA by using calcium phosphate precipitation. At 6 h p.t., medium was removed and fresh complete DMEM containing a final concentration of 500 µM CpK (SiChem; stock solution 100 mM was pre-diluted 1:4 in 1M HEPES shortly before use), and 100 µM ascorbic acid (Sigma Aldrich; stock solution 10 mM) was added. At 44 h p.t., cells were fixed with pre-warmed 2% formaldehyde + 2.5% glutaraldehyde in 0.1 M cacodylate buffer (pH 7.4) for 1.5 h at room temperature, then washed in 0.1 M cacodylate buffer and post-fixed with 2% osmium tetroxide (Electron Microscopy Sciences) for a 1 h on ice. Cells were subsequently dehydrated through an increasing cold ethanol series (30, 50, 70, 80, 90, and 100%; on ice) and two anhydrous acetone series (at room temperature). The coverslip with cells was then removed from the dish, and cells were flat embedded in Epon resin. 70-nm thin sections were cut with an ultramicrotome (Leica EM UC6), collected on formvar-coated 100-mesh copper EM grids (Electron Microscopy Sciences) and stained with a 3% uranyl acetate in 70% MetOH (10 min), and lead citrate (7 min). Cells sections were observed with a JEOL JEM-1400 electron microscope operating at 80 kV (Jeol Ltd.), equipped with a bottom-mounted 4K by 4K pixel digital camera (TemCam F416; TVIPS GmbH).

### CLEM and electron tomography

SupT1 cells were distributed in a 96-well plate (2x10^5^ cells/well; U-bottom; Greiner Bio-one, 650180) and pre-incubated for 16 h with 1 µm aphidicolin (APC; Merck). Cells were pelleted (200 x g, 3 min) and resuspended in complete RPMI medium containing HIV-1*CA14^SIR^ particles (MOI∼0.4). Cells were incubated with viral particles for 120 min at 16°C to adsorb the virus and synchronize virus entry. Samples were then processed for CLEM and ET as described previously (Zila et al., 2021a). In brief, cells were transferred to glass-bottomed microwell of a MatTek dish (MatTek) containing carbon-coated and retronectin-coated sapphire discs (Engineering Office M. Wohlwend). Samples were high pressure frozen, and sapphire discs were then transferred from liquid nitrogen to the freeze-substitution (FS) medium (0.1% uranyl acetate, 2.3% methanol and 2% H_2_O in acetone) tempered at -90°C. Samples were FS-processed and embedded in Lowicryl HM20 resin (Polysciences) according to a modified protocol of Kukulski et al. (Kukulski et al., 2011). For CLEM-ET, thick resin sections (250 nm) were cut and placed on a slot (1 × 2 mm) EM copper grids covered with a formvar film (Electron Microscopy Sciences, FF2010-Cu). Grids were decorated with fiducial marker and stained with Hoechst to visualize nuclear regions. Light microscopy Z stacks of sections were acquired by PerkinElmer UltraVIEW VoX 3D Spinning-disc Confocal Microscope (Perkin Elmer) using a 100 × oil immersion objective (NA 1.49; Nikon), with a *z-*spacing of 200 nm and excitation with the 405-, 488-, 561- and 633-nm laser line. Acquired z stacks were visually examined using Fiji software (Schindelin et al., 2012) and intracellular CA(SiR) positive signals were identified. EM grids were decorated with 15 nm protein-A gold particles for tomogram alignment and stained with uranyl acetate and lead citrate. Grids were loaded to a Tecnai TF20 (FEI) electron microscope (operated at 200 kV) equipped with a field emission gun and a 4K by 4K pixel Eagle CCD camera (FEI). Positions of CA(SiR) signals were pre-correlated with imported SDCM images in SerialEM as described previously (Schorb et al., 2017). Single-axis electron tomograms were carried out. Tomographic tilt ranges were typically from –60° to 60° with an angular increment of 1°. The pixel size was 1.13 nm. Alignments and 3D reconstructions of tomograms were done with IMOD software (Kremer et al., 1996). Post-correlation was performed using eC-CLEM plugin in Icy software (de Chaumont et al., 2012).

### Image analysis

Microscopy images were screened and filtered in Fiji/ImageJ (Schindelin et al., 2012) with a mean filter and background subtraction. Infected cells were quantified in Fiji via segmentation and counting of nuclei and the cell counter to manually quantify the number of positive cells. To determine labeling efficiency of click labeled particles, CA(SiR) intensities of detected immobilized particles based on CA(IF) were quantified using the spot detector of the software Icy (de Chaumont et al., 2012). Five ROIs without particles were measured, and mean intensity in the SiR channel was subtracted as background. The threshold was set to t = 1,000 a.u.. CA(IF) detected spots with intensities above the threshold were classified as CA(SiR) positive.

To analyze particle distribution and intensity measurements throughout the entire volume of cells, z-image series were reconstructed in 3D space using Imaris 9.2 software (Bitplane AG). Individual HIV-1 CA(IF) objects were automatically detected using the spot detector Imaris module, which created for each fluorescent signal a 3D ellipsoid object with 300 nm estimated diameter in *x-y* dimensions and 600 nm in *z*. The local background of each individual spot was subtracted automatically. Subsequently, the mean signal intensity in the CA(SiR) channel was quantitated within all objects. The threshold for SiR intensity was set to t = 7,000 a.u. and adjusted manually for each image by visual inspection. Spots detected in SiR-clusters were excluded. Nuclear objects were manually identified based on the lamin A/C staining. NE-associated objects were classified based on lamin A/C intensities. Every image was manually inspected and a threshold for NE-associated objects was set in the range of 6,300-9,100 a.u.. All other particles were classified as PM/cytoplasm (= in the cytoplasm/at plasma membrane).

To identify post-fusion cores by mCLING ATTO488 staining, CA SiR positive objects were automatically detected using the spot detector Imaris module and the mCLING ATTO488 mean signal intensity co-localizing with each object was quantitated. Particles located outside of the cell, attached to the plasma membrane, or within clusters of foci were excluded from the analysis. Detected objects with the lowest mCLING ATTO488 intensities were individually inspected. Only particles showing a distinct CA SiR focus lacking the mCLING ATTO488 signal were classified as mCLING negative.

Fiji standard ‘greyscale’ lookup table (LUT) was used to visualize single channel images and ‘Fire’ for single channel STED images.

### Data visualization and statistical analysis

Statistical significance was assessed using Prism v9.1.0 (GraphPad Software Inc). A two-tailed non-paired Mann-Whitney test (*α* = 0.05) was used to assess the statistical significance of non-parametric data. Data were plotted using Prism v9.1.0 or the Python statistical data visualization package seaborn v.0.10.0 (Waskom 2020) Graphs show mean/median with error bars as defined in the figure legends.

