## Supplement for "Direct capsid labeling of infectious HIV-1 by genetic code expansion allows detection of largely complete nuclear capsids and suggests nuclear entry of HIV-1 complexes via common routes"

### Supplementary Figures

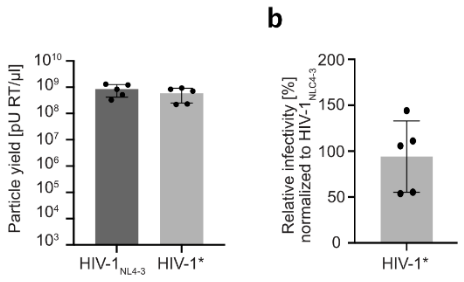


**Figure S1. Characterization of HIV-1*.** The proviral plasmid pNLC4-3 was modified in order to allow for site-specific click labeling of the viral CA. The amber stop codon of the *vpr* open reading frame was mutated into an opal TGA stop codon to avoid GCE modification of Vpr. HEK293T cells were transfected with pNLC4-3 or pNLC4-3* and pNESPylRS-eRF1dn-tRNA and grown in the presence of 500 µM CpK. At 48 h p.t., supernatant of transfected cells was harvested, filtered and concentrated via ultracentrifugation through a sucrose cushion. **(a)** Quantification of RT activity determined in an SG-PERT assay. **(b)** Relative infectivity of HIV-1* measured *via* luciferase assay in TZM-bl reporter cells. Infectivity was normalized to HIV-$1_{NL4-3}$ infectivity measured in parallel. Graphs show mean values and SD of five replicates performed in three independent experiments.

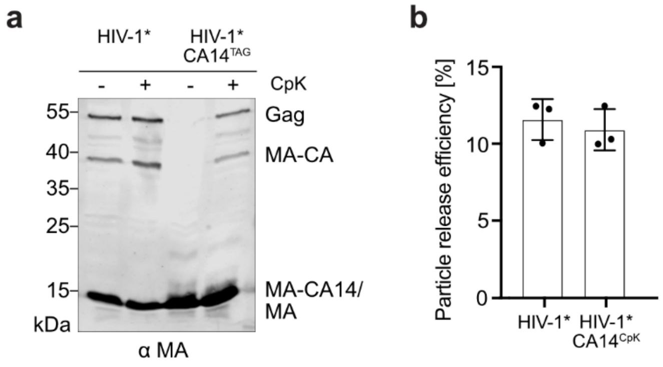


**Figure S2. Detection of full-length Gag Pr55 in the presence or absence of CpK.** Immunoblot analysis of cell lysate of transfected cells. HEK293T cells were co-transfected with pNLC4-3* or pNLC4-3*CA14$^{TAG}$ and pNESPylRS-eRF1dn-tRNA and grown in the presence or absence of 500 µM CpK. **(a)** Cell lysates were separated by SDS-PAGE and proteins were transferred to a nitrocellulose membrane by semi-dry blotting. Gag-derived proteins were detected using polyclonal rabbit antiserum raised against recombinant HIV-1 MA. Note that the truncated MA-CA14 protein (~16,3 kDa) produced in the absence of amber suppression by HIV-1*CA14$^{TAG}$ is not resolved from mature wt MA (~14,8 kDa). Bound antibody was detected using a Li-COR CLx infrared scanner, employing secondary antibody and protocols according to the manufacturer's instructions. **(b)** Supernatant of pNLC4-3* or pNLC4-3*CA14$^{TAG}$ and pNESPylRS-eRF1dn-tRNA transfected HEK293T cells, grown in a 6-well plate and in the presence of 500 µM CpK, was harvested at 48 h p.t.., filtered and concentrated *via* ultracentrifugation through a 20% (w/w) sucrose cushion. Cell and particle lysates were separated by SDS-PAGE and proteins were transferred to a nitrocellulose membrane by semi-dry blotting. Gag derived proteins were detected by quantitative immunoblot (Li-Cor) using a polyclonal rabbit antiserum raised against HIV-1 CA and purified recombinant CA protein as a standard. The ratio of CA amounts detected in the particle lysate to total amounts of Gag derived proteins in cell and particle lysates was calculated to estimate release efficiency for pNLC4-3* (11.6%) and pNLC4-3*CA14$^{TAG}$ (10.9%) transfected cells. Bar graphs represent mean and SD of three technical replicates.

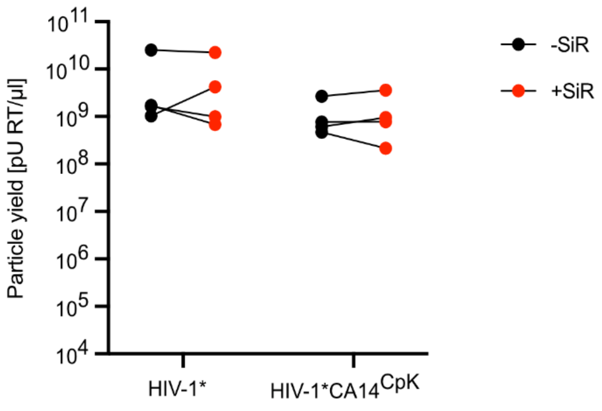

**Figure S3. Effect of click-labeling on particle yield.** HEK293T cells seeded in 6-well plates were transfected with and pNLC4-3* or pNLC4-3*CA14$^{TAG}$ and grown in the presence of 500 µM CpK. Tissue culture supernatant was harvested at 48 h p.i.. One half of the supernatant was incubated with 250 nM SiR-Tet for 30 min, the other half was left untreated. Subsequently, particles were pelleted by ultracentrifugation through a 20% (w/w) sucrose cushion, and pellets were resuspended in 30 µl PBS. The RT activity of unstained (black) and stained (red) samples was determined by SG-PERT. The graph shows data from four parallel transfections; lines connect samples prepared from the same tissue culture supernatant.

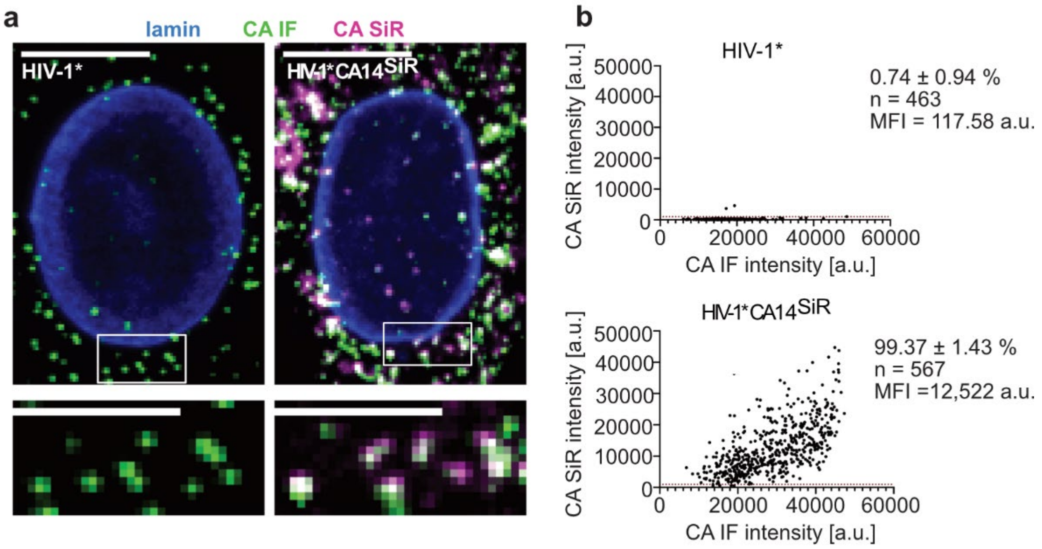


**Figure S4. Detection of click labeled HIV-1*CA14$^{SiR}$ particles in TZM-bl cells.** TZM-bl cells were infected with HIV-1* or HIV-1*CA14$^{SiR}$, treated with 15 µM PF74 for 1 h before fixation at 8 h p.i., immunostaining against lamin A/C (blue) and HIV-1 CA (green) A. and SDCM imaging **(a)** Representative maximum projections from cells infected with HIV-1* or HIV-1* CA14$^{SiR}$. Enlargements of the boxed areas are shown below. Scale bars: 10 µm (cell) and 5 µm (enlargement). **(b)** Mean CA(SiR) intensities plotted against mean CA(IF) intensities of individual intracellular punctae for HIV-1* and HIV-1*CA14$^{SiR}$. The graphs represent data from one of three independent experiments. n=5 cells for HIV-1*CA14$^{SiR}$, n=6 for HIV-1*. The threshold in the SiR channel was set to t = 1,000 a.u. indicated by the red line.

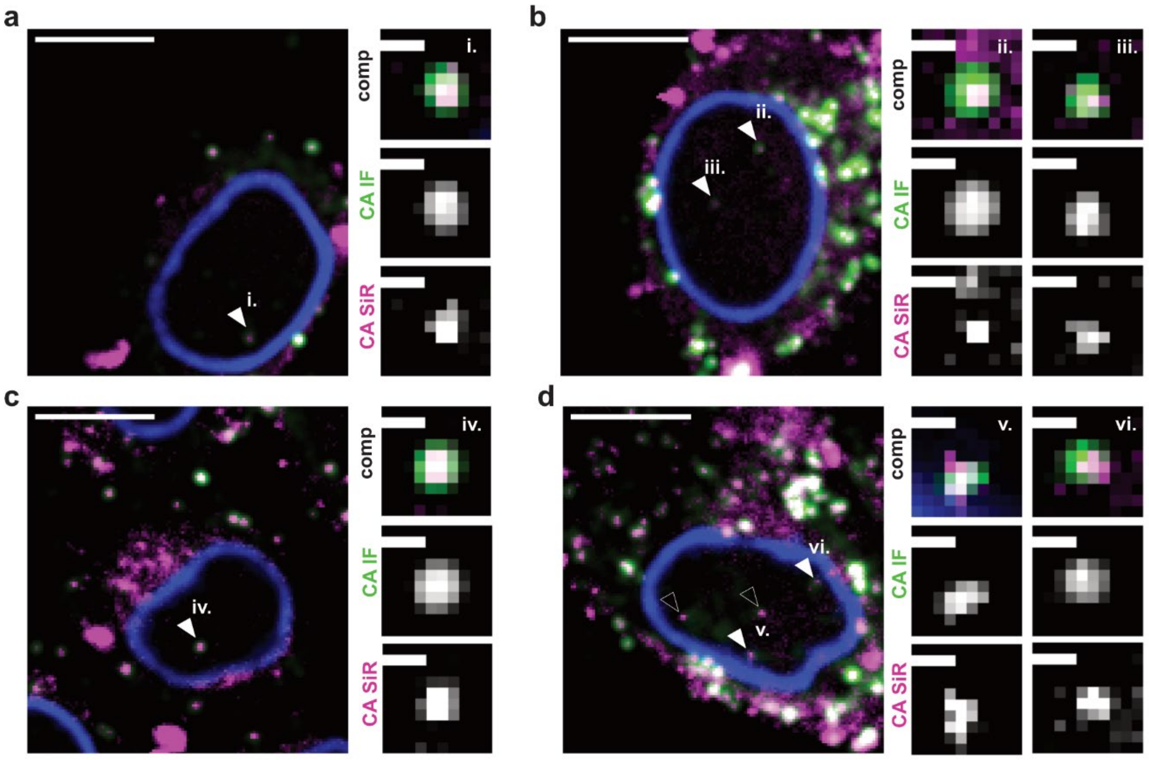


**Figure S5. Confocal micrographs of TZM-bl cells infected with HIV-1*CA14$^{\mathrm{SiR}}$ particles.** Infection was performed using an MOI~0.8 for 18 h p.i.. Cells were fixed, immunostained against lamin A/C (blue) and HIV-1 CA (green) and imaged by SDCM. Images show a single z-slice through the middle of the cells. Arrowheads point towards nuclear CA(IF)/CA(SiR) positive objects (i.-vi.). Enlargements of the indicated nuclear objects are shown to the right of the overview. Mean filter and background subtraction were applied to all images for clarity. Scale bars: 10 µm (overview) and 1 µm (enlargements).

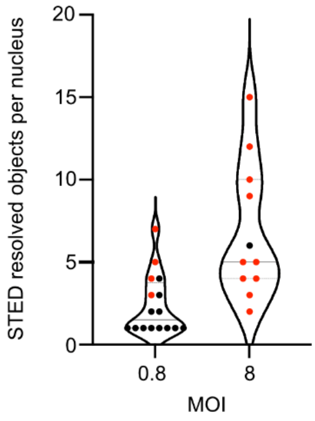


**Figure S6. Number of individual STED resolved particles per nucleus.** TZM-bl cells were infected with HIV-1*CA14$^{SiR}$ at the indicated MOI, treated with 15 µM PF74 for 1h before fixation at 18 h p.i. and immunostained against CA and lamin A/C as described in materials and methods. CA(IF) was imaged in confocal mode, CA(SiR) in confocal and STED mode. Cells displaying at least one diffraction limited CA signal within the nucleus were analyzed in STED mode to determine the total number of individual particles. Dots represent individual cells (MOI 0.8: mean = 2.4, n = 16; MOI 8: mean = 6.8, n = 11). Black: exclusively individual particles detected; red: cell comprises at least one cluster. Graphs show median and interquartile lines in grey.

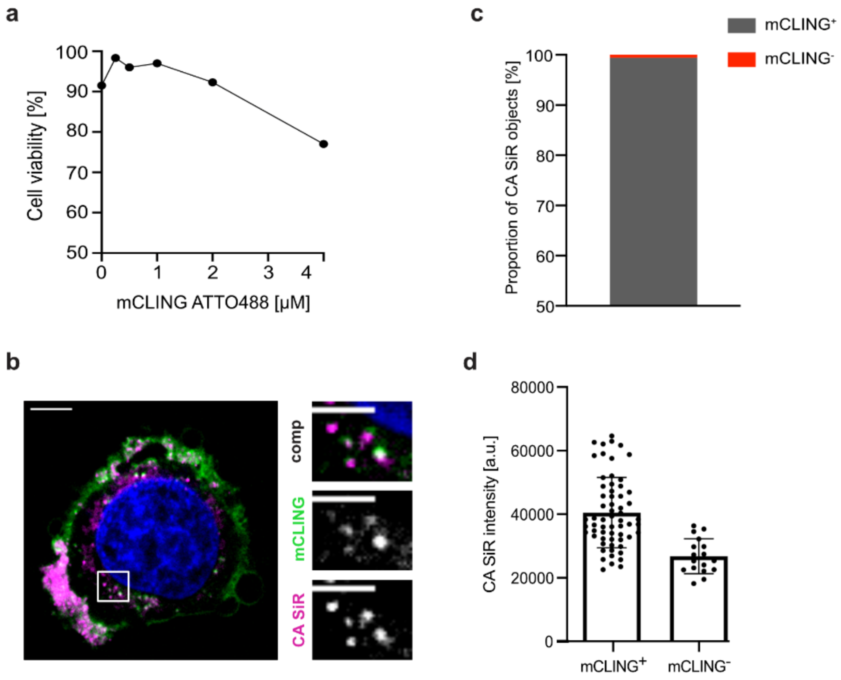

**Figure S7. The majority of HIV-1*CA14$^{SiR}$ particles in the cytosol of HeLa derived cells is detected in endosomal vesicles. (a)** To test viability of cells in in the presence of mCLING ATTO488, the compound was titrated on TZM-bl cells. Cells were incubated for 30 min at 16°C and before counting using an automated cell counter (Greiner Bio-one). Data represent the mean of one experiment performed in duplicate. **(b, c)** Following incubation with 2 µM mCLING ATTO488 at 16°C for 30 min, TZM-bl cells were infected with 10 µU RT/cell at 37°C, fixed at 3 h p.i. with 4% PFA + 0.2% GA and imaged by SDCM. **(b)** Representative image from one experiment (single z-slice through the middle of the cell). Enlargement of the boxed area of the cell (right) shows colocalization of mCLING ATTO488 (green) and CA SiR (magenta) signals. Scale bars: 10 µm (overview) and 5 µm (enlargement). **(c)** Quantitation of mCLING ATTO488 positive (98.78 %) and negative (1.22 %) CA SiR objects (total n = 2,606, 5 cells) as described in materials and methods. **(d)** Quantitation of CA SiR signals of mCLING ATTO488 positive (n=62, mean=41,397 ± 10,909 a.u.) and negative (n=17, mean=27,655 ± 5,812 a.u.) objects in the cytosol. Graphs represent mean values and SD.

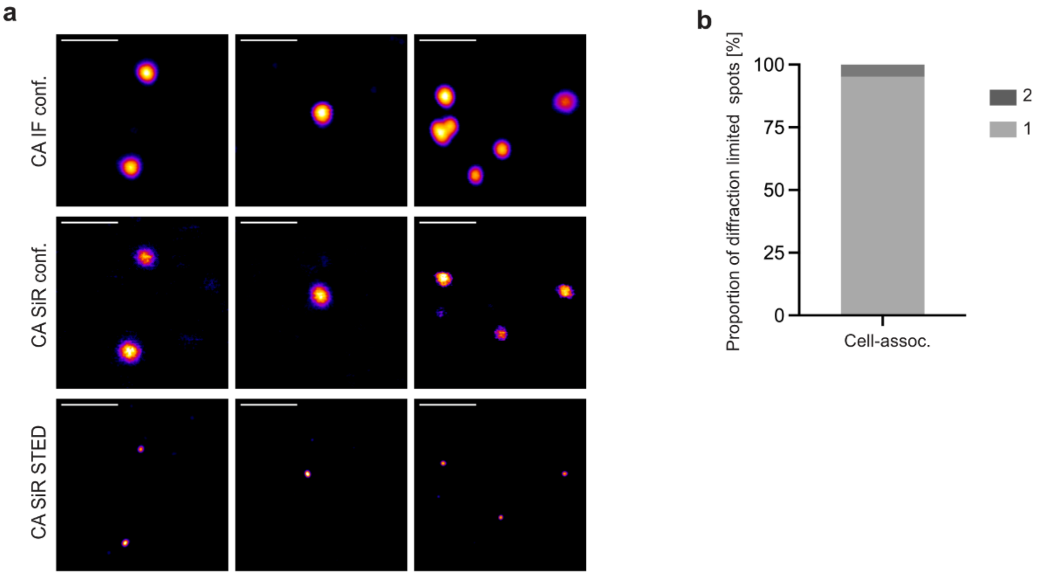


**Figure S8. STED nanoscopy of CA(IF)/CA(SiR) double positive objects in the cytoplasm of TZM-bl cells. (a)** TZM-bl cells were infected with HIV-1*CA14$^{SiR}$ (MOI~0.8) for 18 h, fixed and immunostained against HIV-1 CA, and imaged using an Abberior STED system. Cytoplasmic objects were detected by CA(IF) (top panel) and CA(SiR) (middle panel). Diffraction limited double positive foci localized in confocal mode were imaged in STED mode in the SiR channel (bottom panel). Mean filter and background subtraction was applied to all images for clarity. Scale bars: 1 µm. **(b)** The number of individual capsids per diffraction limited spot (n=83) was determined from STED images. ~95% (n=79) of analyzed objects corresponded to an individual particle, while ~5% (n=4) of foci were resolved into two objects by nanoscopy.

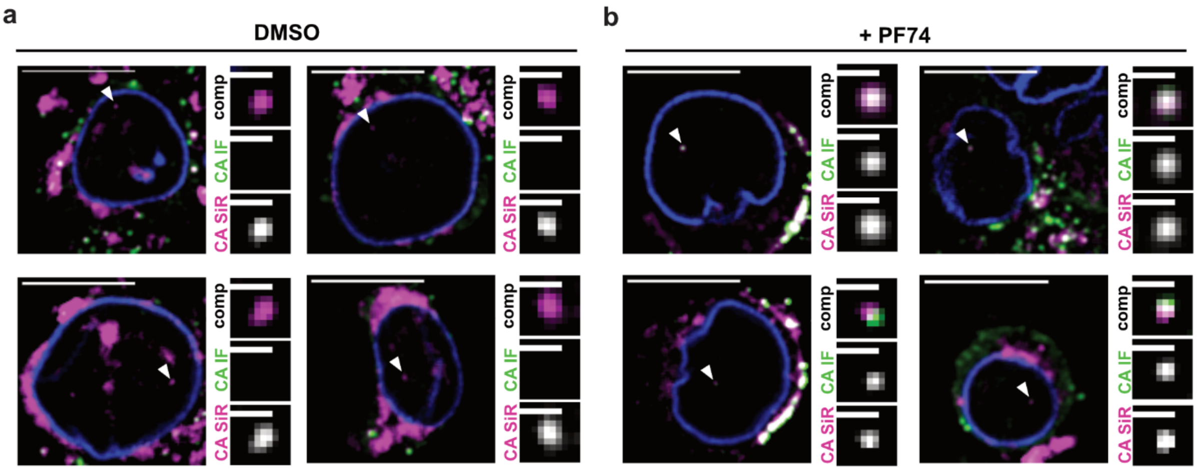


**Figure S9. Confocal micrographs of primary CD4⁺ T cells infected with HIV-1*CA14$^{SiR}$.** Cells were infected (MOI~0.8, determined in TZM-bl cells) and subsequently treated with DMSO **(a)** or 15 µM PF74 **(b)** for 1 h before fixation at 24 h p.i., immunostaining against CA (green) and lamin B1 (blue) and SDCM. Images show a single z-slice through the middle of the cell; two representative examples are shown per condition. Arrowheads point towards nuclear CA(IF)/CA(SiR) double positive objects. Enlargements of the indicated complexes are shown on the right. Mean filter and background subtraction was applied on each image for clarity. Scale bars: 10 µm (overview) and 1 µm (enlargements).

**Supplementary Movie 1: CLEM-ET analysis of nuclear HIV-1*CA14$^{SiR}$ capsid-like structures in T cells.** Related to Figure 5e and 5f. Shown is a segmented and isosurface rendered tomographic reconstruction to highlight the morphology of several clustered capsid-related structures visualized inside the nucleus of an infected SupT1 T cell (upon APC treatment), correlated with the position of a CA(SiR) signal.
